# Combining antigenic data from public sources gives an early indication of the immune escape of emerging virus variants

**DOI:** 10.1101/2021.12.31.474032

**Authors:** Antonia Netzl, Sina Türeli, Eric B. LeGresley, Barbara Mühlemann, Samuel H. Wilks, Derek J. Smith

## Abstract

The rapid spread of the Omicron BA.1 (B.1.1.529.1) SARS-CoV-2 (Severe Acute Respiratory Syndrome Coronavirus 2) variant in 2021 resulted in international efforts to quickly assess its escape from immunity generated by vaccines and previous infections. Numerous laboratories published BA.1 neutralization data as preprints and reports. We collated this data in real time and regularly presented updates of the aggregated results in US, European and WHO research and advisory settings. Here, we retrospectively analyzed the accuracy of these aggregations from 85 different sources published during a time period from 2021/12/08 up to 2022/08/14. We found that the mean titer fold change from wild type-like variants to BA.1, a standard measure of a variant’s immune escape, remained stable after the first 15 days of data reporting in people who were twice vaccinated, and incoming data increased the confidence in this quantity. Further, it is possible to build reliable, stable antigenic maps from this collated data already after one month of incoming data. We here demonstrate that combining early reports from variable, independent sources can rapidly indicate a new virus variant’s immune escape and can therefore be of immense benefit for public health.

## 1 Introduction

The WHO classified the Omicron BA.1 variant (B.1.1.529+BA.1) as a Variant of Concern (VoC) on November 26, 2021[1], and BA.1 quickly replaced Delta as the world-wide dominant variant. Since then, SARS-CoV-2 continues evolving and other variants took over. The rapid emergence and replacement of the dominant SARS-CoV-2 variant required quick reactions and global public health efforts to assess its immune escape in different contexts of vaccination and infection history. At the time of BA.1’s emergence, multiple laboratories rapidly produced virus neutralization data with diverse serum and variant panels and released them quickly as preprints or prelimnary reports for public use, before journal publication. To aggregate the information from individual sources for research and policy guidance, we incrementally collected and summarized available data in openly accessible documents between December 2021 and August 2022 [2, 3]. This open and collaborative approach to science in response to the COVID-19 (Coronavirus disease-19) pandemic was crucial for informed public health policies. Here, we retrospectively analyze the data we collated in real time with a focus on how quickly summary results of BA.1’s immune escape reach a level of confidence relevant for public health guidance.

In late 2021, population immunity against SARS-CoV-2 in Europe and the US consisted of people infected with Wuhan-1/D614G, Alpha, Beta or Delta, and a large proportion of recipients of the two dose Wu-1 (Wuhan-1) vaccines. Wu-1 vaccines performed exceptionally well against the early SARS-CoV-2 variants, showing that up to that point the antigenic evolution of SARS-CoV-2 was moderate and did not necessitate a vaccine strain update [4]. BA.1 started to circulate around the time the recommendation for a third vaccine, or 1^st^ booster dose, was issued [5, 6]. Its rapid takeover and infection of Wu-1 vaccinated people raised concerns about the protection conferred by an additional Wu-1 vaccine dose [7].

Reacting quickly to the emergence of a virus variant is essential to keep infections, hospitalizations and deaths low. Such reactions include the assessment of the novel variant’s properties, including its transmissibility, severity, and potential to escape population immunity to inform public health decisions, such as advising for a vaccine strain update or implementing non-pharmaceutical interventions. Ideally, data to inform such decisions should be from controlled, reliable trials with large numbers of, preferably randomized, subjects. However, this type of data usually takes considerable time to generate which impedes a quick emergency response. With regards to the assessment of BA.1’s ability to escape population immunity, independent research and public health laboratories generated data quickly, albeit with smaller sample sizes and at times substantial variations across laboratories due to assay type, virus isolates, and type of and time since exposure of sera. After BA.1’s emergence, we responded to the quick action by the multiple individual laboratories and collated the not-yet peer reviewed data that was published on preprint servers in real time. Here, we show that the limitation of small sample sizes and variation of data from independent sources can be overcome when combining data from multiple independent laboratories, giving a rapid indication of a variant’s escape potential to further public health decisions. Incremental versions of these data and analyses [2] were frequently presented in national (UK and US), regional (WHO euro) and global (WHO) scientific and public health fora as part of the scientific and public health response to BA.1 [8–10]. Now, using publicly available data from 2021/12/08 up to 2022/08/14, we show that collated data from different sources can be used to produce stable vaccine escape measures soon after variant emergence, providing valuable information for a quick response to emergencies.

## 2 Results

### 2.1 Data collection

We analyzed Omicron BA.1 neutralization geometric mean titers (GMT) from 85 different sources which at the time of data collection (2021/12/08–2022/08/14) were mainly in not peer-reviewed preprint form on bioRxiv, medRxiv or otherwise in the public domain. By now, many of these preprints have been published in peer-reviewed journals. However, we collected the data in real time and updated a publicly available Google Slide deck [2], summarising each study, and a publicly accessible google sheet document incrementally with incoming data [3]. We base our analysis here on such collected data, the first publicly available preprints, in order to emulate the real-world scenario of a novel emerging variant and the urgency that comes with reporting its mmune escape.

The collected data include neutralization of various Omicron sublineages (B.1.1.529: BA.1 and BA.1+R346K (BA.1.1); BA.2, BA.2.12.1 and BA.2.75; BA.3; BA.4/5) as well as ancestral and other SARS-CoV-2 variants by different vaccine sera and sera of individuals infected with the ancestral virus (614D/G, from here onwards referred to as wild type WT), Alpha (B.1.1.7), Beta (B.1.351), Gamma (P.1) or Delta (B.1.617.2) variant. As the data was generated in different laboratories with little coordination between laboratories, a variety of neutralization assays and cell types was used, an overview is given in Table 1. We categorized the serum panels used by the different laboratories by their infection or vaccination history into different serum groups as described in the methods section.

**Table 1:**
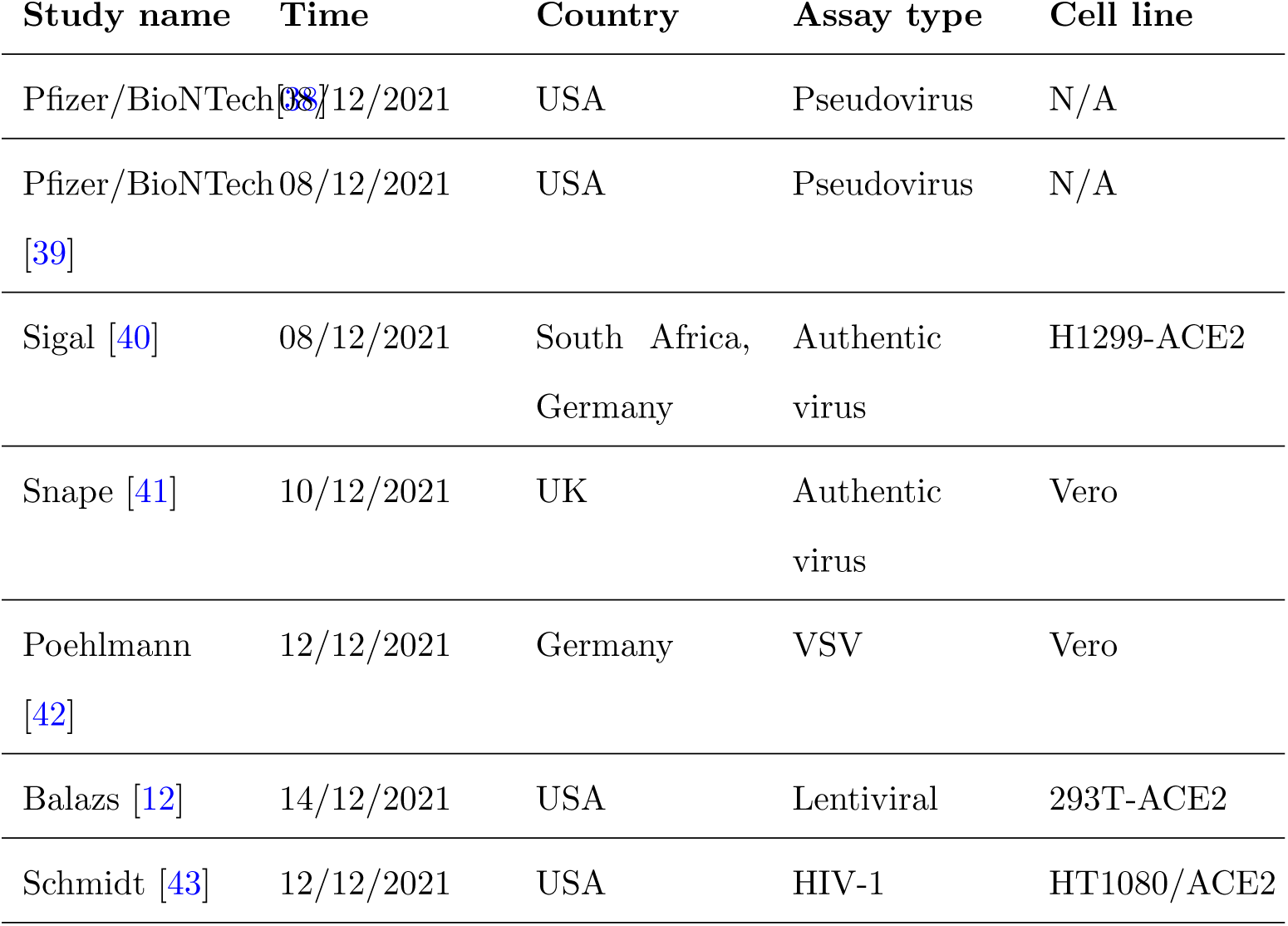
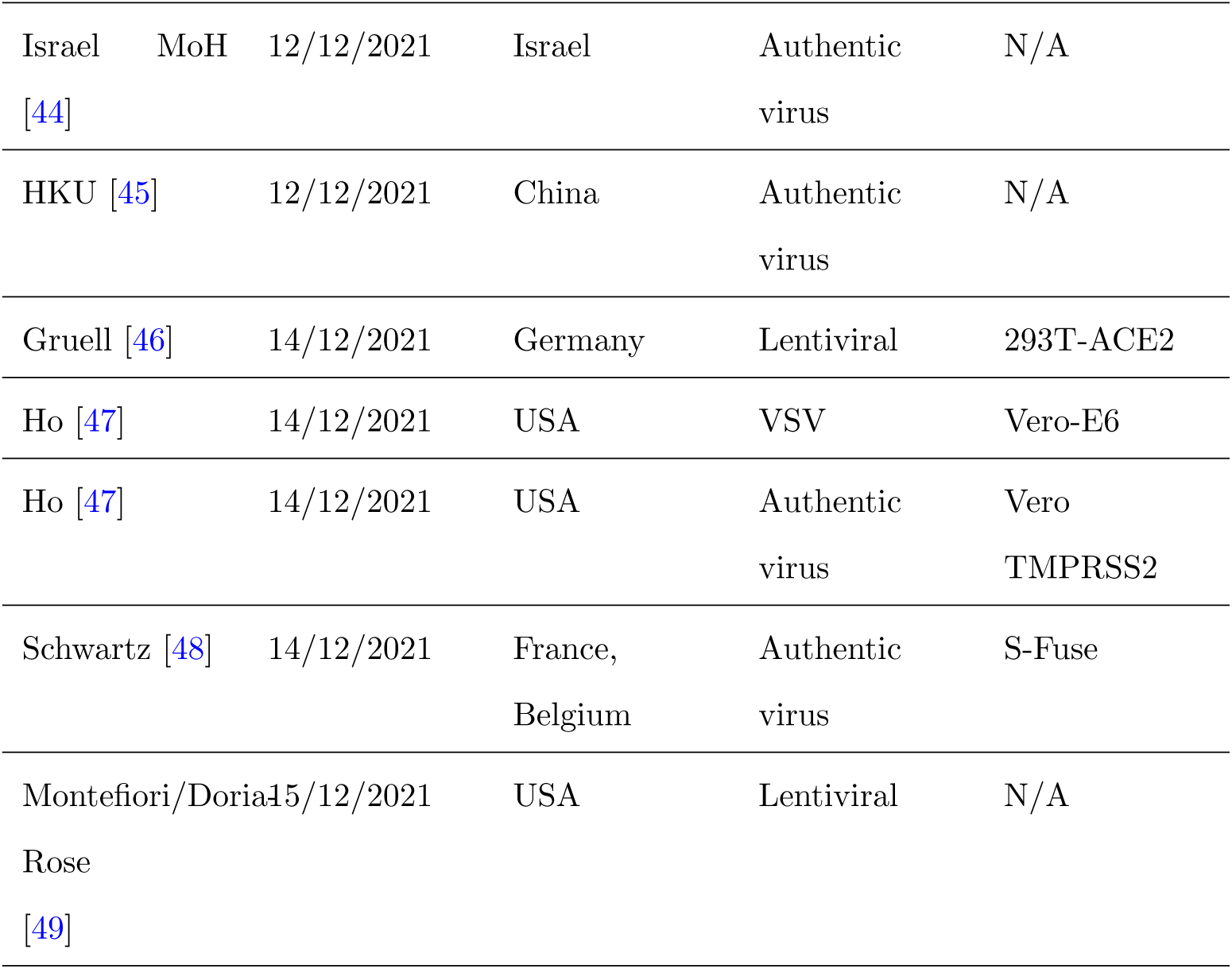
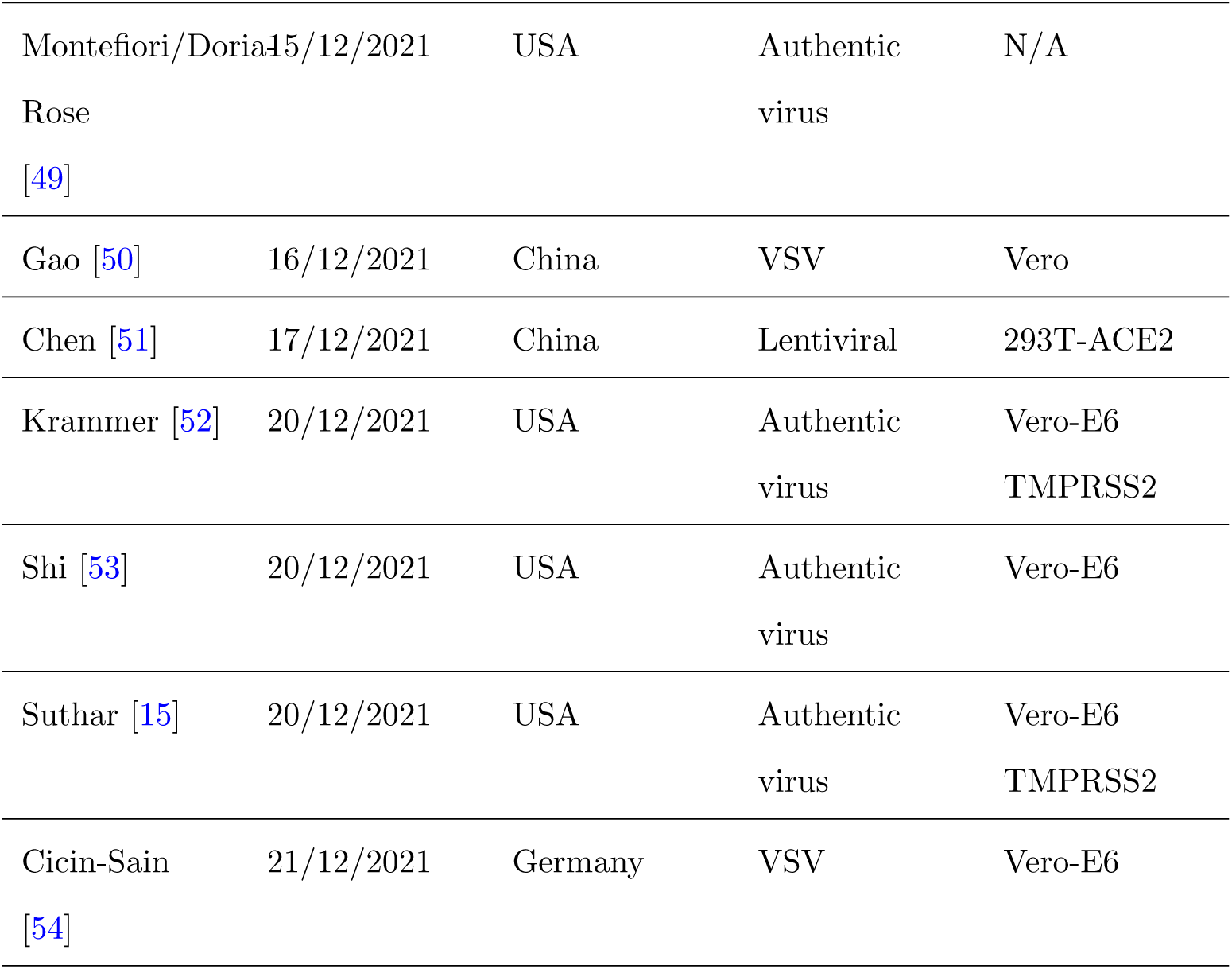
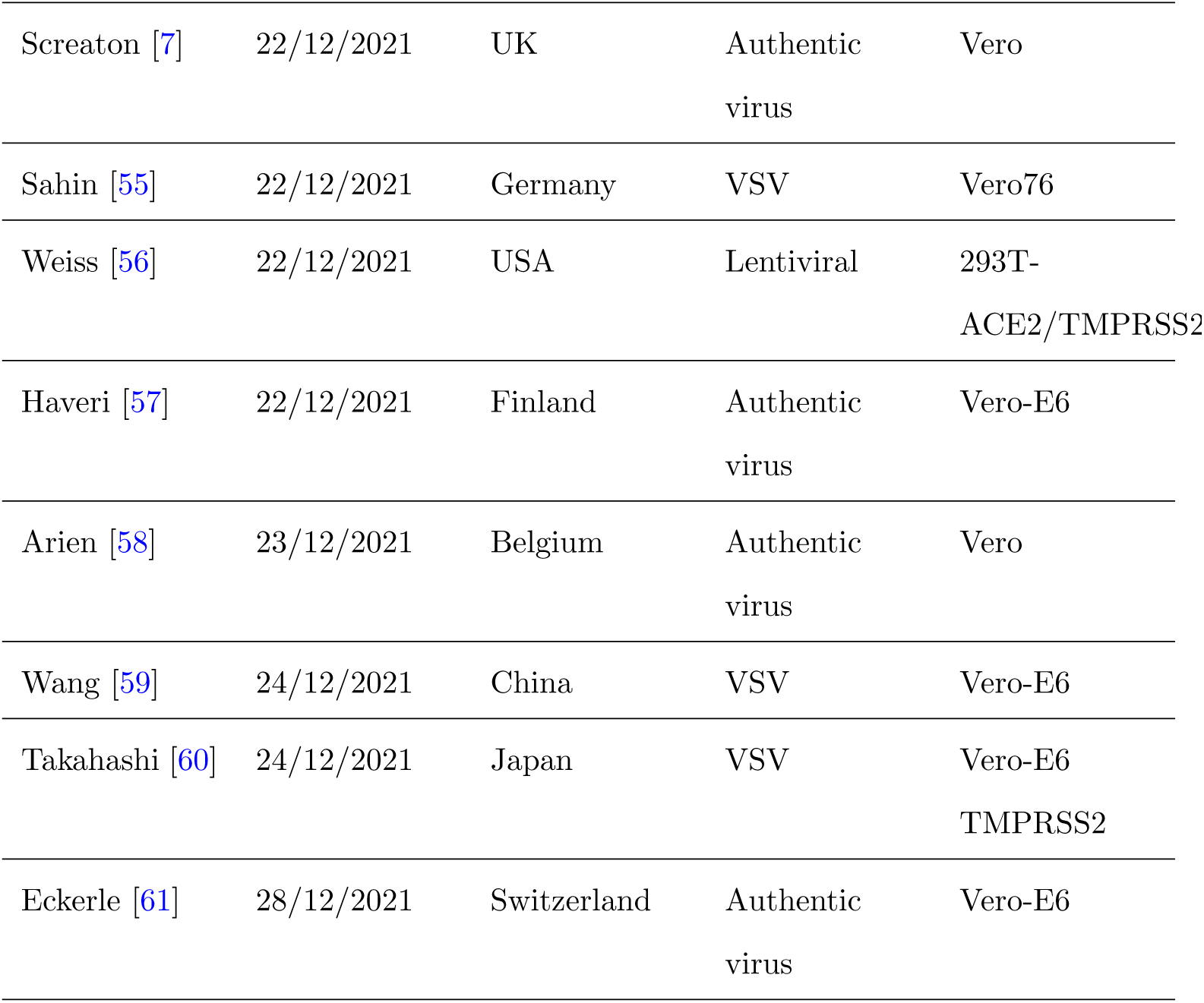
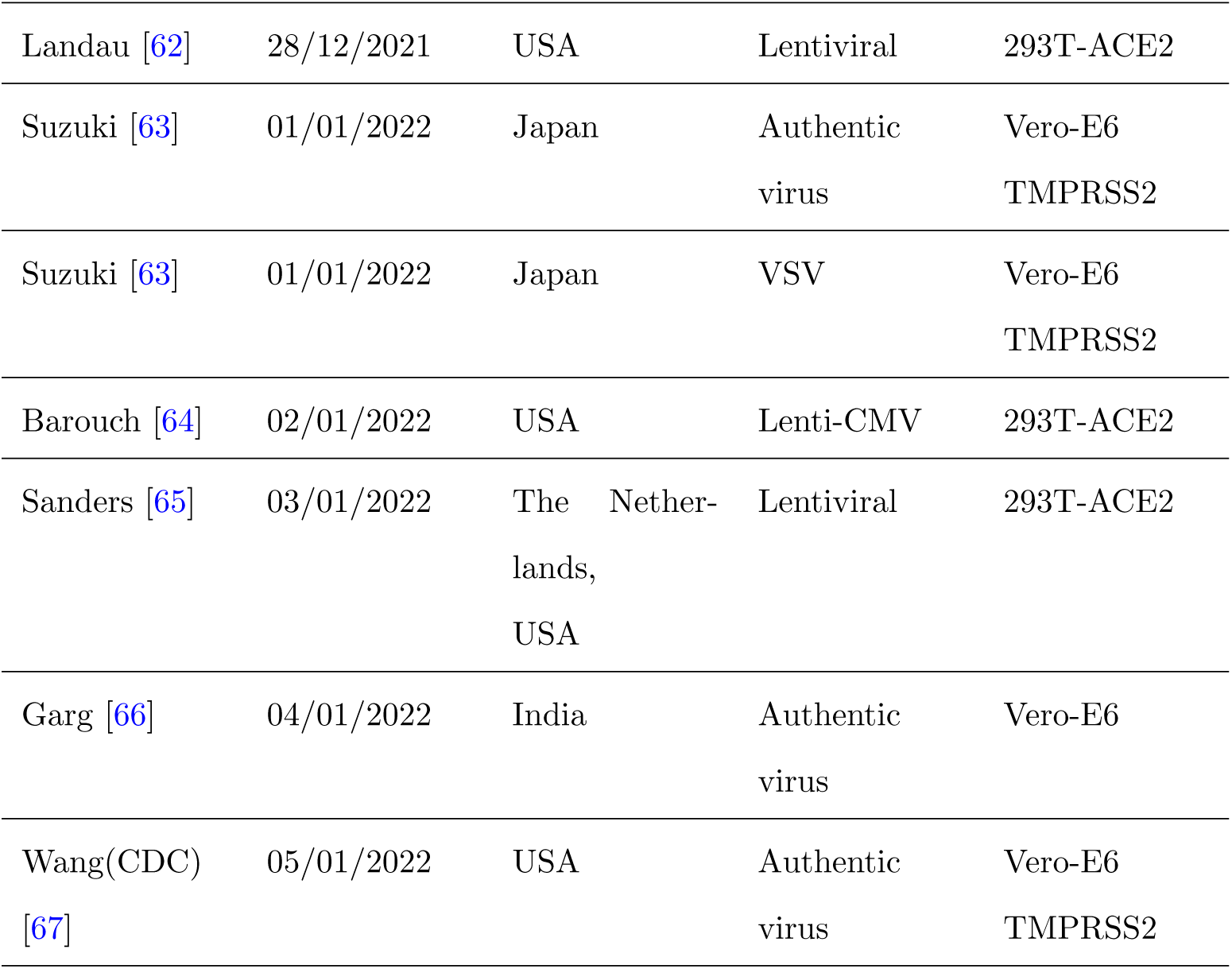
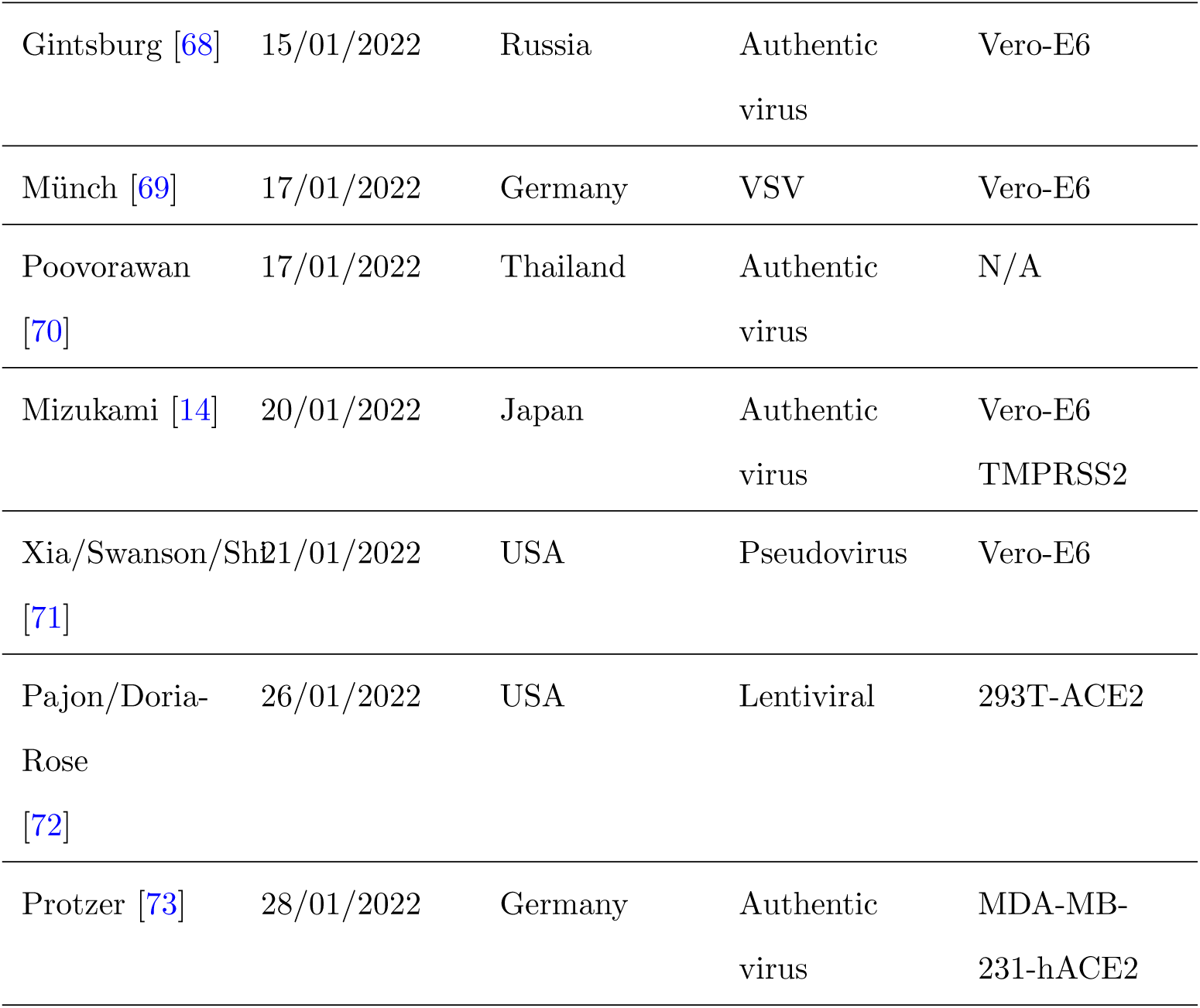
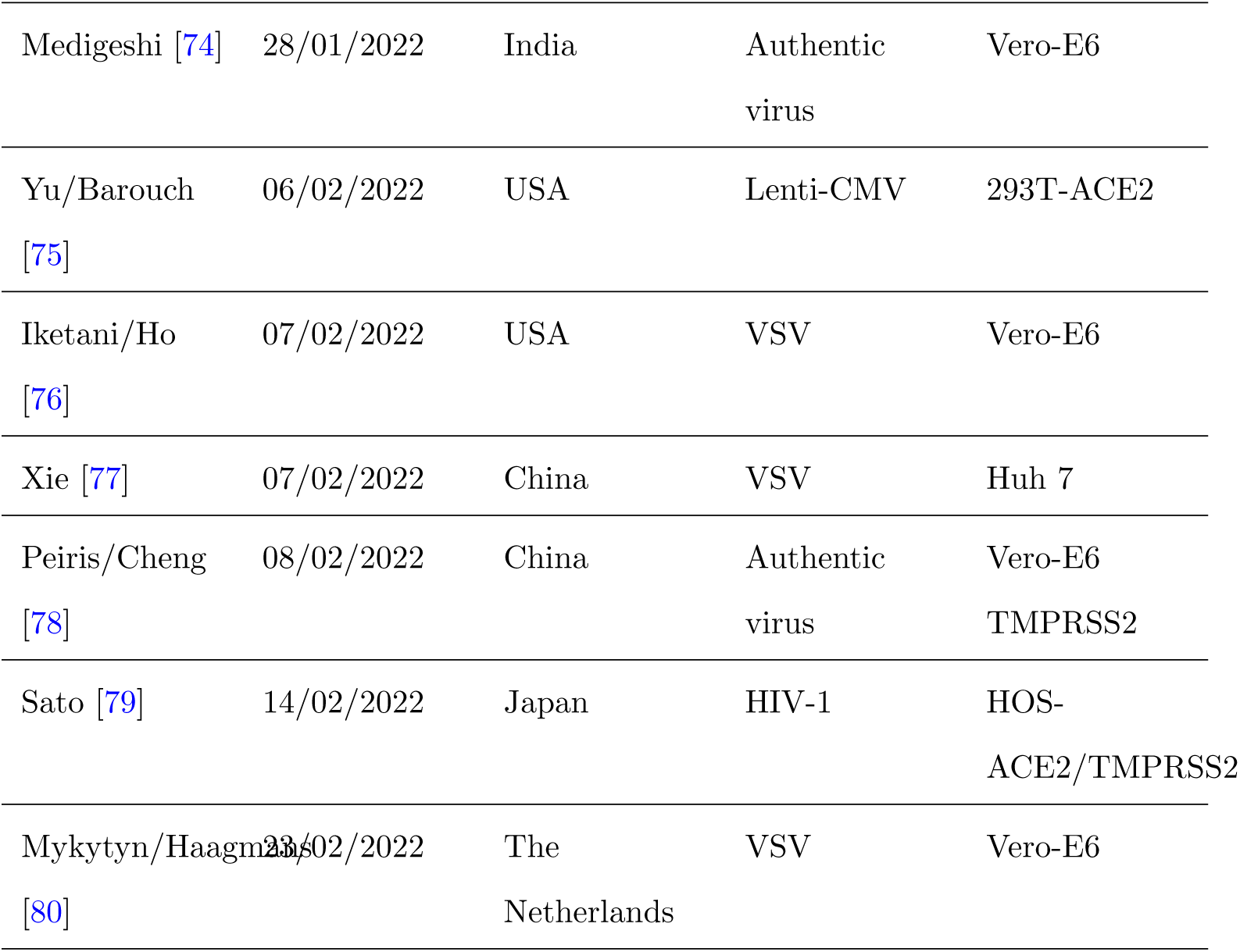
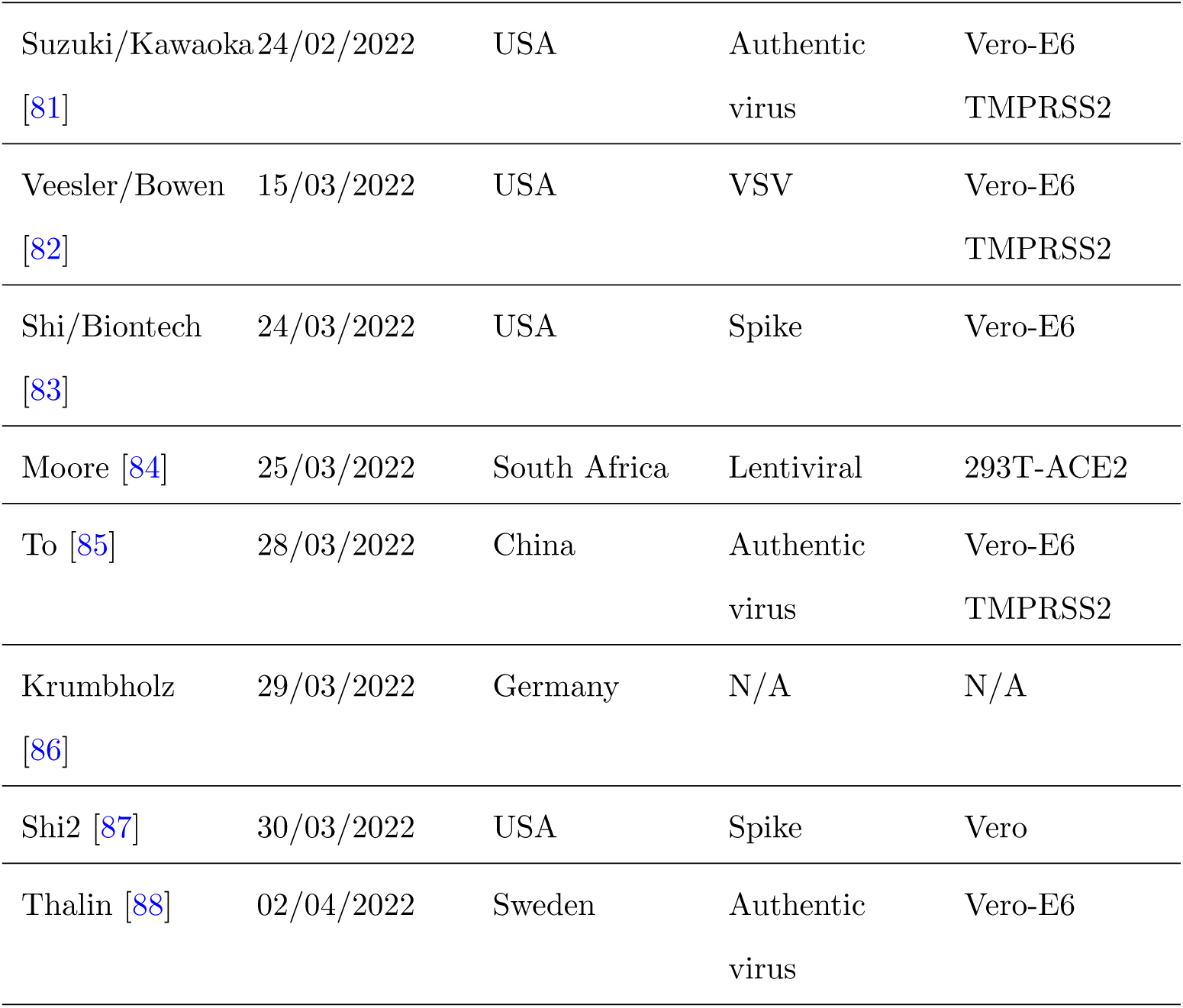
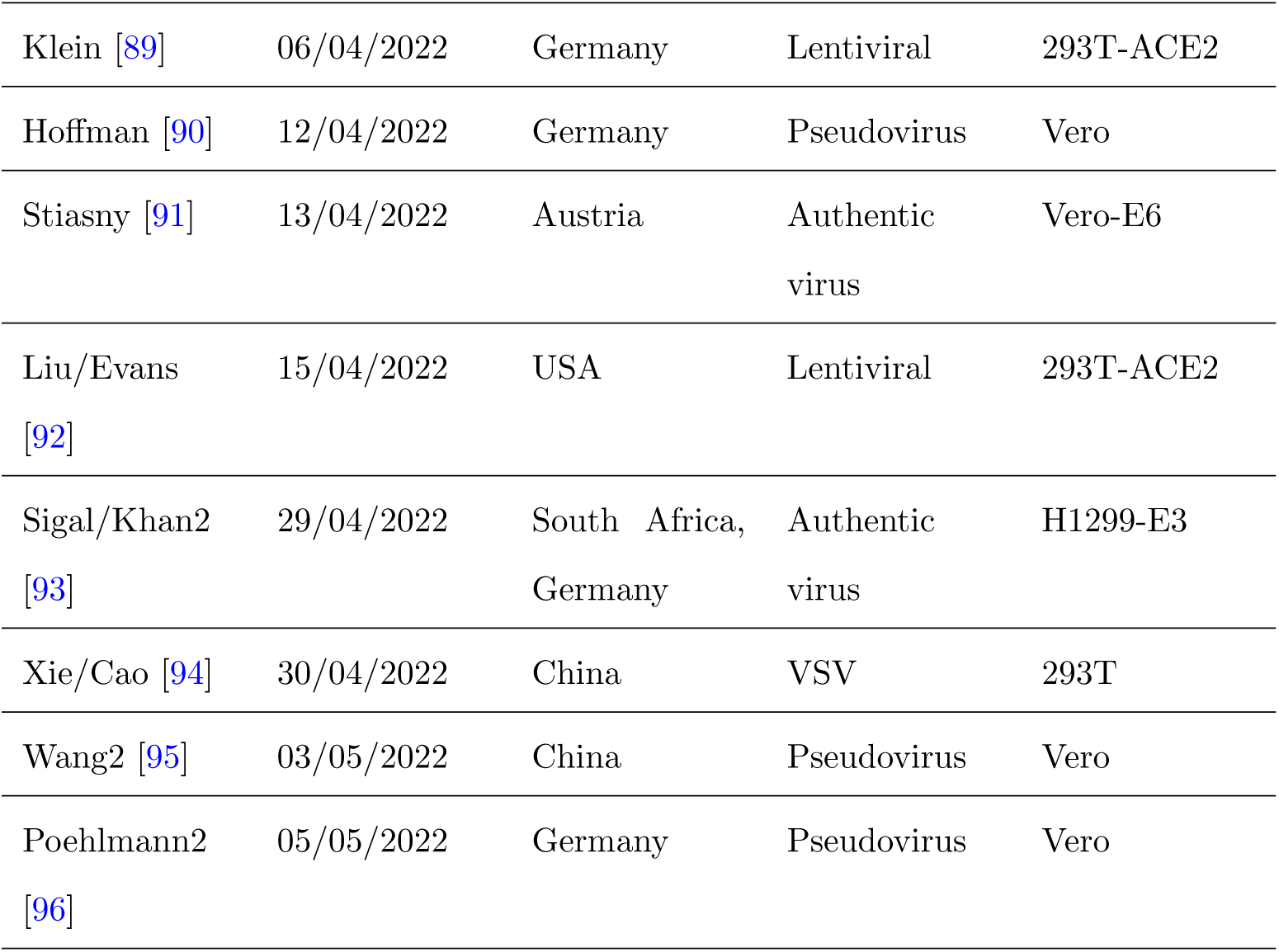
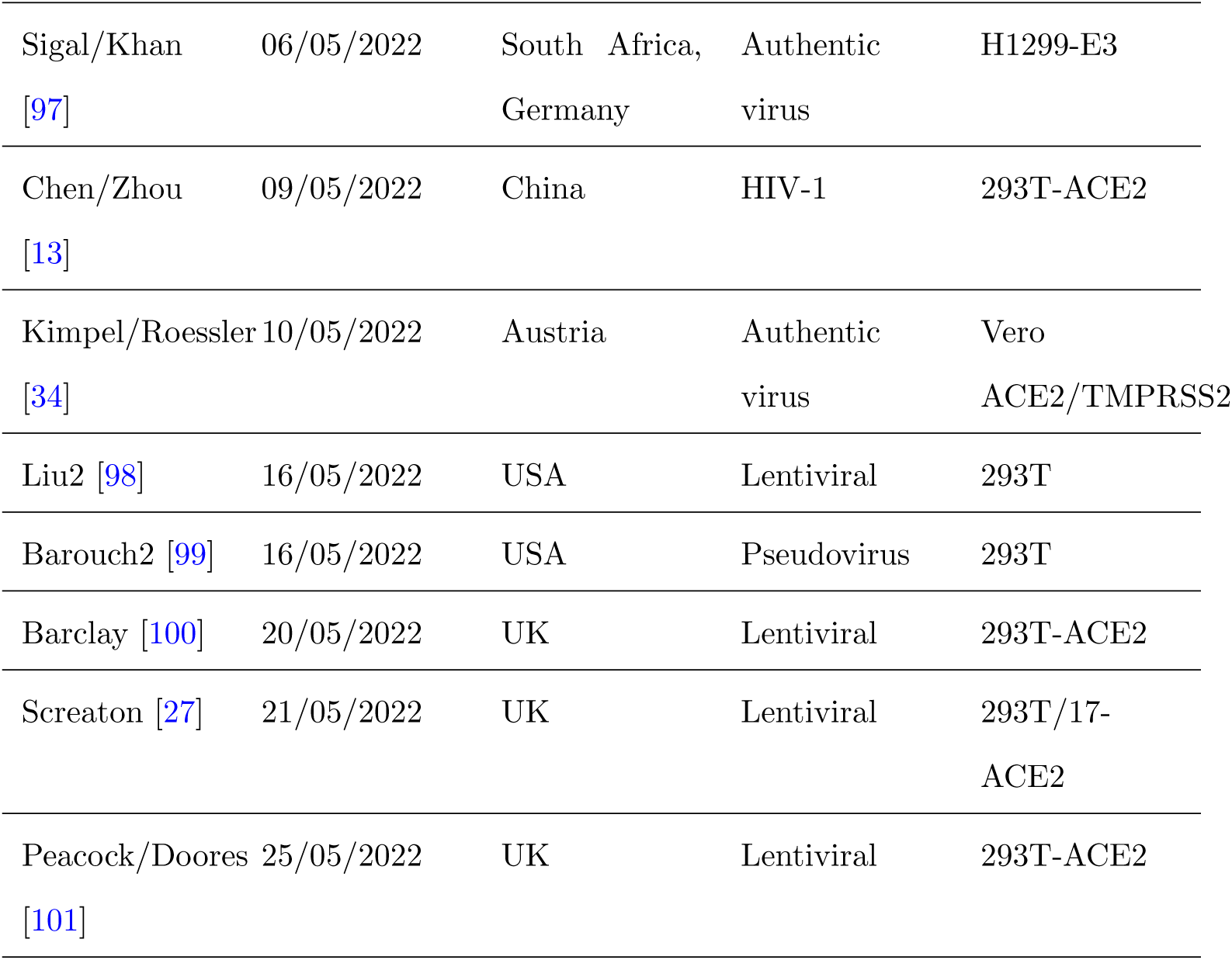
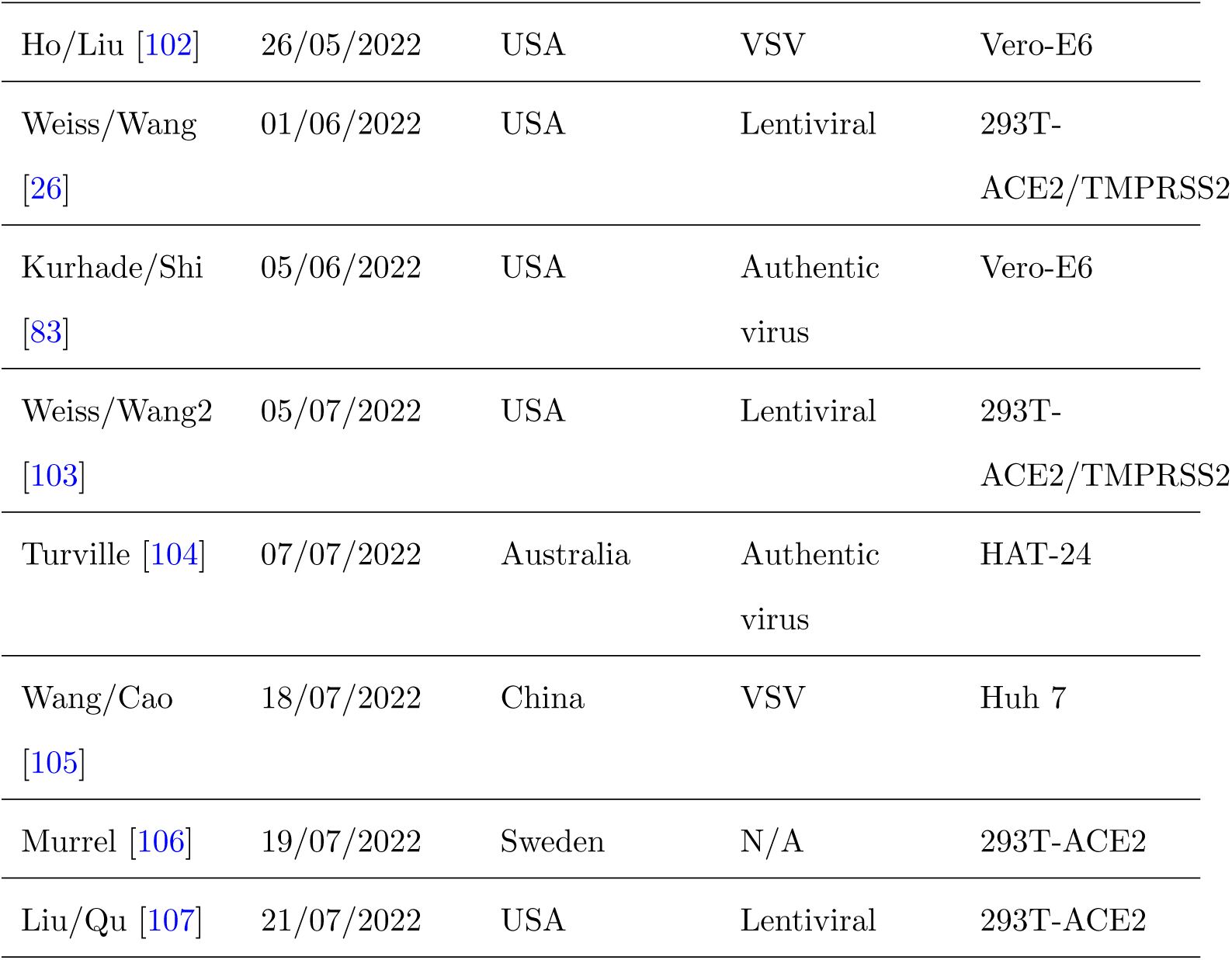
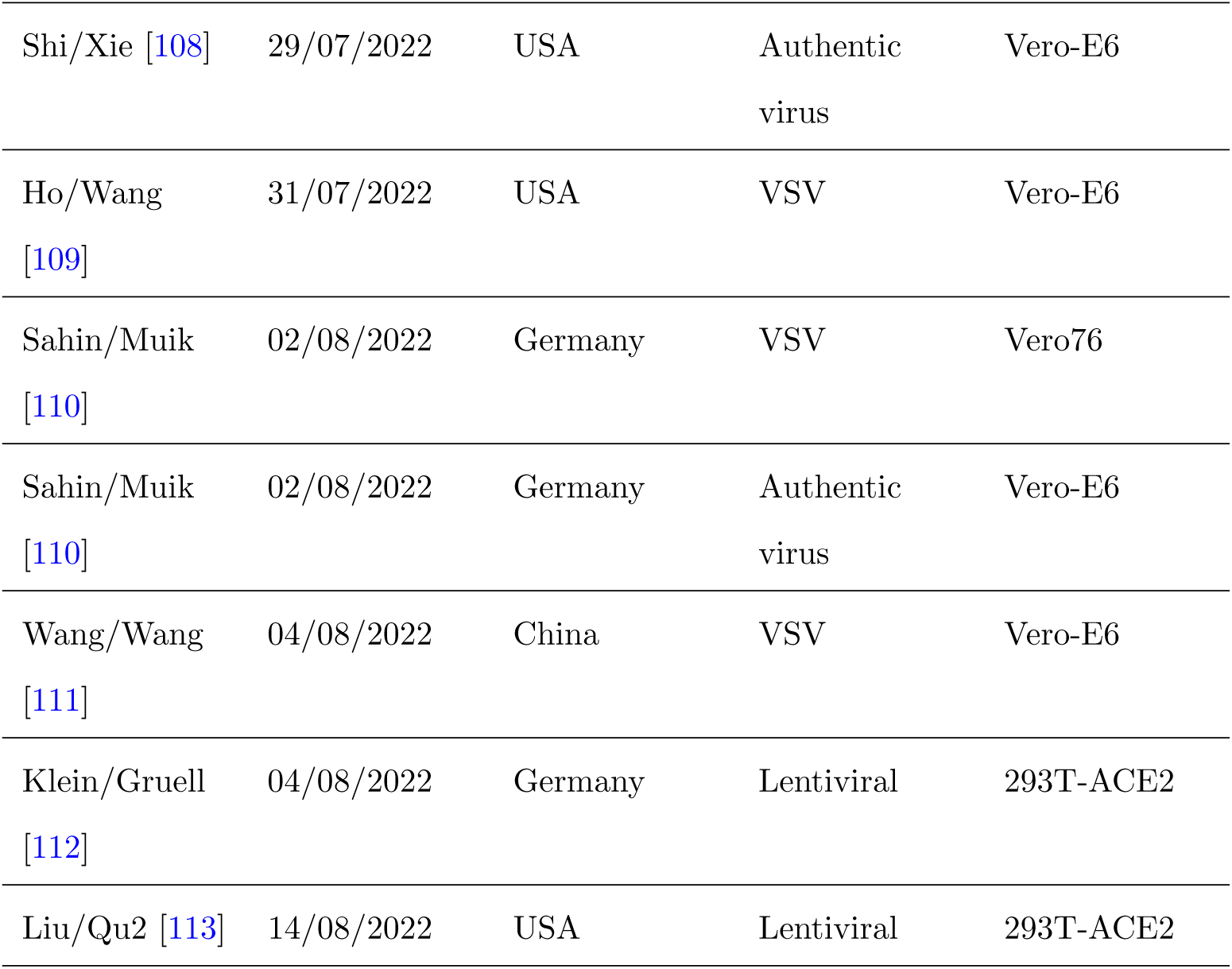
List of included studies. An overview of the collected studies. The link to the document from where the data was extracted is given in [3]. The time lists the upload date of the study extracted from the source link, or when the data was added by the authors (DD/MM/YYYY). The studies were named after the first or the corresponding author, the corresponding author’s country affiliation is listed in the Country column. The neutralization assay virus type is given in the assay type column and refers to a specific pseudotype if not labelled “Pseudovirus” or “Authentic virus”. The cell line shows which cells were used in the neutralization assay. “N/A” indicates that no information was found

The majority of the collected data was generated using the Omicron BA.1 lineage. Some research groups indicated that the virus they used had the R346K mutation (BA.1.1, BA.1+R346K). To identify whether this substitution impacted neutralization, we compared BA.1 and BA.1.1 neutralization titers in the same sera and found GMTs to be highly consistent for the two lineages (Supplementary Fig. A1, Supplementary Table A1), therefore we did not differentiate between the two in our downstream analysis. As time progressed, other Omicron lineages emerged and we kept collecting the neutralization data as it was generated. However, for the purpose of this study we focused our analysis on the BA.1 lineage.

### 2.2 Collective data can inform public health decisions early after variant emergence

To inform public health decisions, such as vaccine strain updates, after the emergence of a new virus variant, immune escape measures like fold change of titers to a vaccine variant are used. Fold change of neutralization titers are a standard measurement of immune escape and are used as input for the antigenic characterization of viruses [11]. Data from government approved laboratories and clinical trials using standardized assays contribute hugely to informing these decisions but take time to generate. At the same time, individual studies generate these data quickly, but in a less controlled manner. To assess BA.1’s immune escape from WT, we extracted the reported mean fold changes or calculated fold changes from reported geometric mean titers (GMTs) of individual studies in different serum cohorts. We did this incrementally, adding new data in real time and summarizing it in public documents [2], and we show data within the first two weeks of reporting in Fig. 1a.

**Fig. 1.**
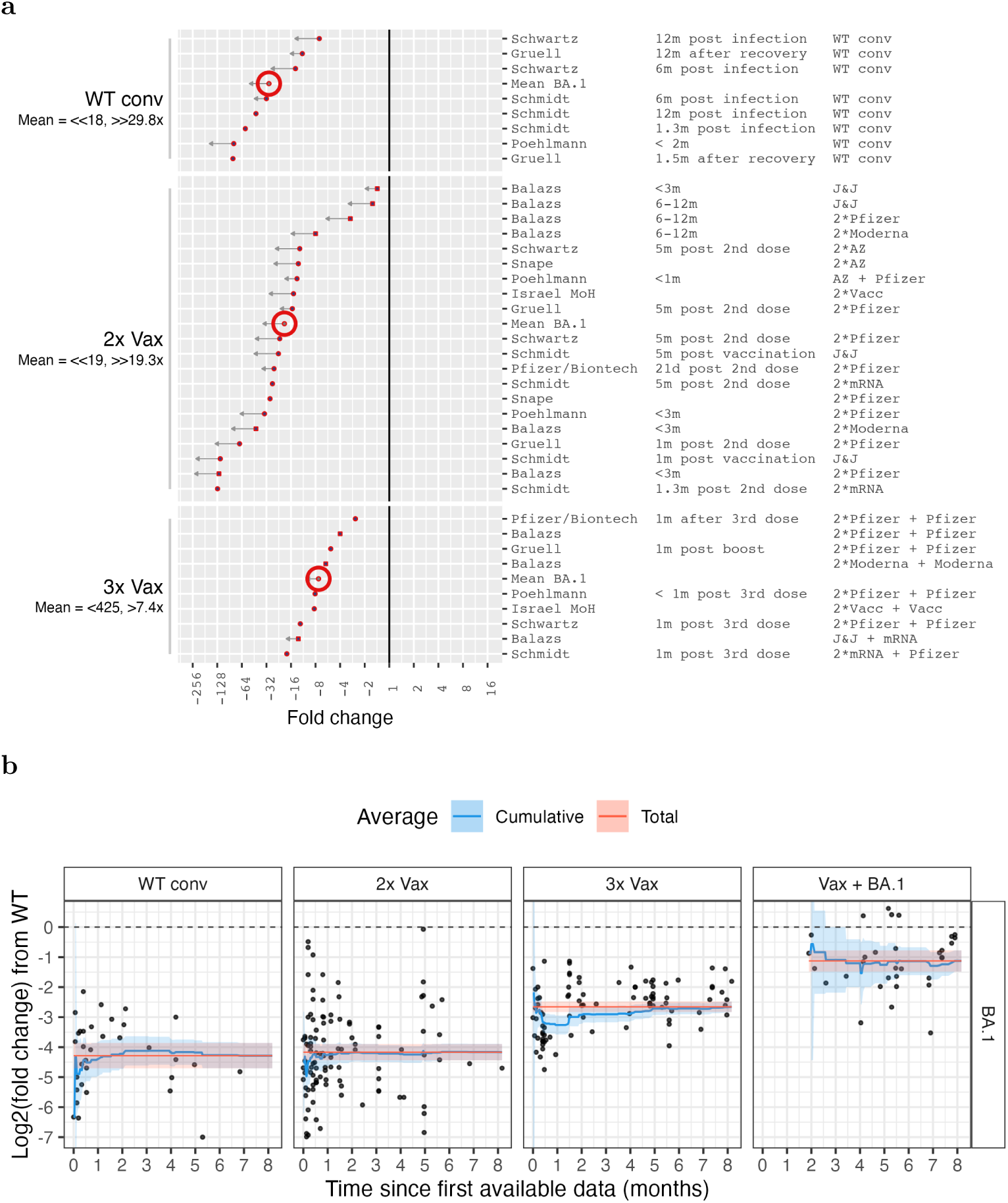
BA.1 neutralization titer fold change from WT (614D/G) titers over time. **a)** Fold changes of GMT neutralization titers as reported by individual studies are shown for the first two weeks of data reporting, ordered by magnitude and stratified by serum group. Row labels show the study name (Table 1), time since exposure and serum type. The big red circle indicates the mean value for each group. Below the serum group label, the BA.1 geometric mean titer (GMT) is given followed by the mean fold change from WT titers. Arrows in the plot and “*<*/*>*” in the label indicate uncertainties in the point estimate due to titers below the limit of detection (LOD) of the assay. A short arrow (“*>*”/”*<*”) marks measurements with more than half of BA.1 titers below the assay’s LOD, or conversely reference antigen titers at or lower than the LOD. Long arrows (“*>>*”/”*<<*”) mark measurements with more than approximately 80% of BA.1 titers below the LOD. The solid vertical line marks no fold change. **b)** Log2 fold changes from WT GMTs to BA.1 GMTs are shown for individual studies by time since the first reported data (1 month = 30 days). The blue line shows the cumulative mean with 95%CI indicated59by the shaded area, the red line with shaded area corresponds to the mean fold change with 95%CI at the end of the data collection period. The data is stratified by exposure history

We continued collating data and were interested how soon after variant emergence enough data was available to obtain stable immune escape estimates relevant for public health guidance. To do so, we examined how the mean neutralization titer fold drop from WT to BA.1 changes over time with the addition of more data (Fig. 1b). For WT conv (614D/G convalescent) and 2x Vax (twice vaccinated), there was already enough data reported within the first 15 days of data reporting such that the cumulative mean was within the 95%CI (confidence interval) of the total mean and remained stable over the remaining collection period. In the 3x Vax group, the fold change from WT was overestimated by one two-fold from two weeks onwards and approached the overall mean thereonafter.

We calculated the confidence intervals of the cumulative means as a measure of how reliable the mean estimates are up to a given time point. We found that 2 weeks of data collection for the WT conv, 2x Vax and 3x Vax group was sufficient to bring the cumulative mean %95CI of all three groups to a single log2 unit precision. After a month of data collection it was further improved up to half a log2 unit for the Vax groups. The cumulative mean %95CI for the Vax + BA.1 group was generally wider throughout data collection, especially early on, owing to less data availability for this type of sera. Nevertheless, the %95CI for the Vax + BA.1 was less than two log2 units one month after the first Vax + BA.1 data was collected, and less than one and a half og2 units after a month and a half.

In addition, we found clear evidence of the benefit of a third vaccine dose over two doses already after two weeks, which only became more certain over time. Further, we found that an exposure to BA.1 after vaccination greatly reduces BA.1’s immune escape. This observation was only possible once enough BA.1 breakthrough infections occurred to be assayed. Despite this lag in reporting, employing public data indicated the benefit of a vaccine strain update to BA.1 already 4 months after its emergence. These observations are further explored in section “Antigenic cartography of BA.1’s mmune escape” via use of antibody landscapes.

To get a confidence estimate of the variance in the titer fold change calculation after say ten studies we randomly selected (with replacement) ten of all studies of 2x Vax data. We repeated this random selection (bootstrapping) 11 times, corresponding to 10% of the number of studies with 2x Vax data, and calculated the mean WT to BA.1 fold change each time. We next calculated the 95%CI of this distribution of mean fold changes and repeated this bootstrap process for all number of studies from 1 to 108, and additionally for the 3x Vax data (Supplementary Fig. 2). We found that with just 10 randomly selected studies, the overall fold change was within the 95%CI of the mean fold changes for both the 2x and 3x Vax cohorts. With 25 and 15 random studies, respectively, the range of lower to upper 95%CI bound was lower than 1.5x, meaning that both would be within one dilution step of a neutralization assay.

### 2.3 Omicron BA.1’s immune escape from wild type in different exposure histories

In the same style as Fig. 1a, we present the BA.1 fold drops from WT at the final stage of data collection in Fig. 2 for more types of exposure histories and more labs for exposure groups that already exist in Fig. 1a. The numerical data in all serum groups and for other variants is summarized in Tables 2-5.

**Fig. 2.**
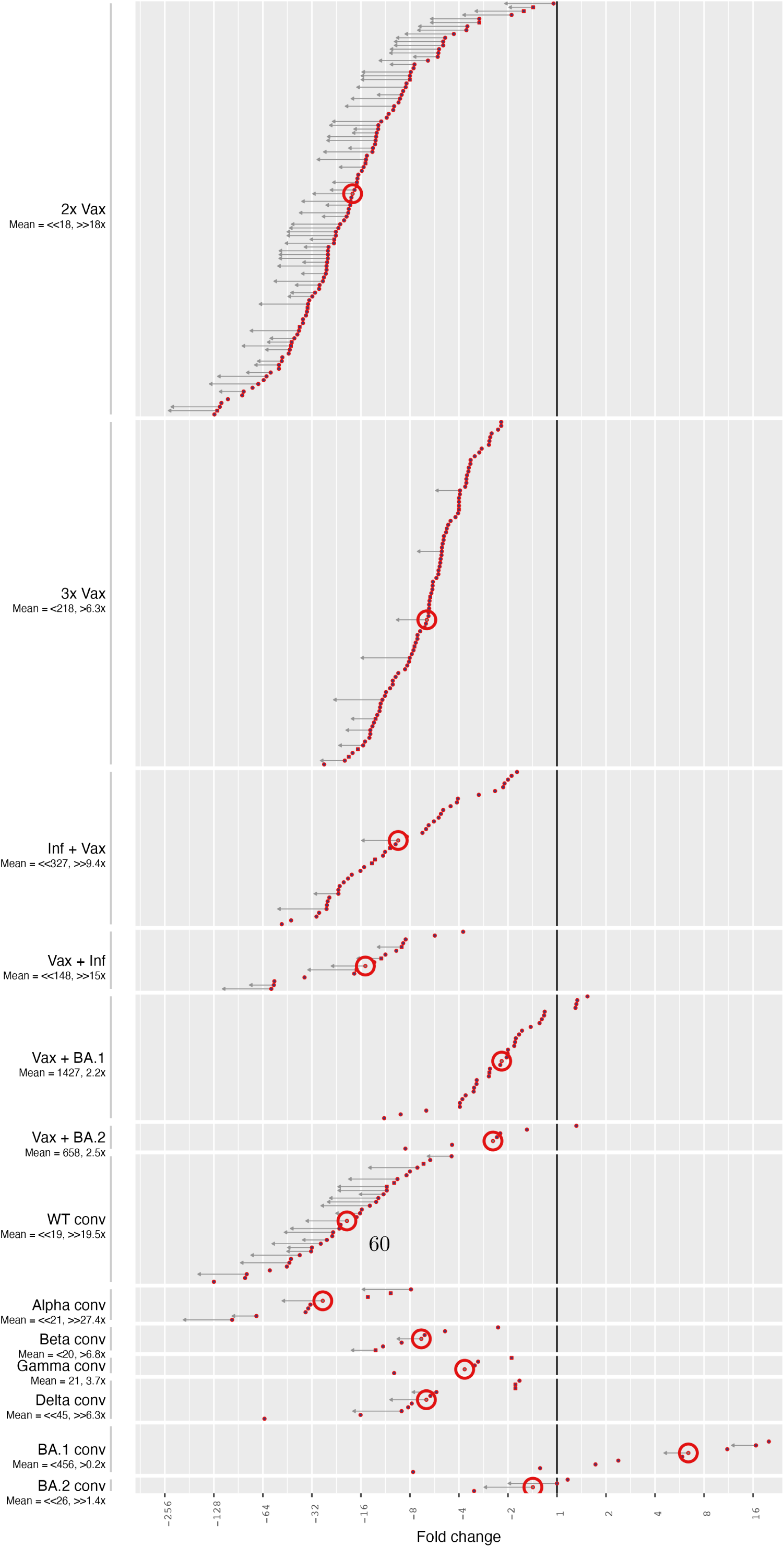
Omicron BA.1 neutralization titer fold changes relative to WT (614D/G). Same as Fig. 1a but without row labels

**Table 2:** Variant Geometric Mean Titers (GMT) per serum group (conv = convalescent). The 95%CI is given in parentheses, the number of data points n in the next line. WT summarizes wild type-like strains (e.g: 614D, 614G)

| Serum Group | WT | Alpha | Beta | Gamma | Delta | BA.1 |
| --- | --- | --- | --- | --- | --- | --- |
| 2x Vax | 334 (253; 440)<br>n=87 | 173 (70; 426)<br>n=10 | 75 (53; 105)<br>n=40 | 153 (43; 541)<br>n=6 | 123 (81; 185)<br>n=44 | 18 (15; 22)<br>n=87 |
| 3x Vax | 1344 (1014; 1781)<br>n=82 | 453 (200; 1026)<br>n=9 | 354 (210; 598)<br>n=27 |  | 398 (245; 648)<br>n=32 | 221 (163; 300)<br>n=82 |
| Inf + Vax | 2840 (1621; 4977)<br>n=31 | 683<br>n=1 | 661 (250; 1743)<br>n=9 | 360<br>n=1 | 1677 (851; 3305)<br>n=16 | 348 (182; 665)<br>n=31 |
| Vax + Inf | 3353 (1281; 8775)<br>n=10 | 2223 (294; 16813)<br>n=2 | 1090 (34; 34774)<br>n=3 | 1112<br>n=1 | 2100 (761; 5792)<br>n=10 | 190 (51; 708)<br>n=10 |
| Vax + BA.1 | 3272 (2039; 5250)<br>n=31 | 807 (120; 5434)<br>n=5 | 1120 (427; 2942)<br>n=10 |  | 942 (467; 1903)<br>n=17 | 1505 (977; 2319)<br>n=31 |
| Vax + BA.2 | 1626 (699; 3782)<br>n=6 | 1486 (1302; 1697)<br>n=2 | 547 (377; 793)<br>n=2 |  | 668 (265; 1683)<br>n=3 | 658 (193; 2240)<br>n=6 |
| WT conv | 405 (246; 667)<br>n=29 | 176 (30; 1044)<br>n=6 | 56 (17; 184)<br>n=7 | 37 (2; 565)<br>n=3 | 183 (89; 376)<br>n=16 | 18 (13; 27)<br>n=29 |

Table 2: **Variant Geometric Mean Titers (GMT) per serum group (conv = convalescent).** The 95%CI is given in parentheses, the number of data points n in the next line. WT summarizes wild type-like strains (e.g: 614D, 614G) (Continued)
| Serum Group | WT | Alpha | Beta | Gamma | Delta | BA.1 |
| --- | --- | --- | --- | --- | --- | --- |
| Alpha conv | 668 (65; 6815)<br>n=6 | 699 (58; 8501)<br>n=5 | 142 (12; 1679)<br>n=5 | 66 (0; 29728)<br>n=3 | 328 (23; 4763)<br>n=6 | 23 (7; 84)<br>n=6 |
| Beta conv | 110 (9; 1290)<br>n=5 | 133 (2; 10303)<br>n=4 | 298 (11; 7976)<br>n=4 | 158 (0; 596407)<br>n=3 | 75 (10; 589)<br>n=5 | 27 (3; 215)<br>n=5 |
| Gamma conv | 67 (4; 1086)<br>n=3 | 77 (2; 3341)<br>n=3 | 103 (2; 5508)<br>n=3 | 275 (0; 179386178094)<br>n=2 | 26 (0; 1470)<br>n=3 | 20 (1; 466)<br>n=3 |
| Delta conv | 437 (81; 2371)<br>n=8 | 195 (14; 2706)<br>n=5 | 72 (5; 1090)<br>n=5 | 70 (3; 1939)<br>n=3 | 2294 (451; 11673)<br>n=8 | 50 (14; 185)<br>n=8 |
| BA.1 conv | 71 (26; 194)<br>n=12 | 40 (3; 460)<br>n=4 | 62 (19; 205)<br>n=7 | 16 (0; 31041577)<br>n=2 | 53 (17; 167)<br>n=10 | 456 (144; 1438)<br>n=12 |
| BA.2 conv | 37 (0; 6933)<br>n=3 | 11<br>n=1 | 11<br>n=1 | 5<br>n=1 | 15 (0; 3013)<br>n=2 | 26 (1; 840)<br>n=3 |

The double and triple vaccinated serum groups constituted the majority of the data that have been reported and consequently analyzed here and were the most relevant from a public health perspective at the time of data collection. The 2x Vax serum group contained the highest number of individual measurements and exhibited the widest spread and largest uncertainty in fold drops of BA.1 neutralization compared to WT. The 2x Vax group showed the most variability in fold changes from WT, which we attribute to a wide range of reported WT titers, serum collection times from two weeks to nine months post second dose, and limit of detection (LOD) censoring of low to non-detectable Omicron titers (Supplementary Fig. A3-A4). We found an average fold drop of 18x in this serum group (Fig. 1, Table 3) when treating measurements below an assay’s LOD in the common manner as LOD/2. However, the majority of fold changes were likely greater than the point estimates due to many BA.1 titers being below the LOD. Consequently, the average fold drop is likely substantially greater than 18x. We found that in many studies with an unexpectedly low fold drop from WT to BA.1 titers, titers against WT were very low [12–15]. Low titers against the reference antigen limit the amount of further reduction until an assay’s detection imit is reached, resulting in LOD censoring of titers and seemingly low fold drops (Supplementary Fig. A3).

**Table 3:** BA.1 titer fold changes from variants per serum group (conv = convalescent). Mean fold drops were calculated from fold drops per study, not from GMT fold drops. Studies reporting fold drops but not GMTs result in a discrepancy between GMT based mean fold drop and individual study based mean fold drop (Table 2). The 95%CI is given in parentheses, the number of data points n in the next line. WT summarizes wild type-like strains (e.g: 614D, 614G)

| Serum Group | WT | Alpha | Beta | Gamma | Delta |
| --- | --- | --- | --- | --- | --- |
| 2x Vax | 18 (14.9; 21.7)<br>n=108 | 14.6 (6.3; 34.2)<br>n=11 | 4.2 (3.3; 5.3)<br>n=44 | 18.2 (8.4; 39.8)<br>n=7 | 6.8 (5.2; 8.8)<br>n=51 |
| 3x Vax | 6.3 (5.6; 7.1)<br>n=90 | 4 (3.4; 4.6)<br>n=9 | 2.6 (2.1; 3.1)<br>n=30 |  | 2.9 (2.4; 3.5)<br>n=34 |
| Inf + Vax | 9.4 (7; 12.8)<br>n=40 | 10.5<br>n=1 | 5.9 (3.7; 9.3)<br>n=12 | 5.5<br>n=1 | 4.7 (2.5; 8.9)<br>n=16 |
| Vax + Inf | 15 (9.4; 24)<br>n=15 | 15.1 (0.9; 242.4)<br>n=2 | 3 (1.7; 5.4)<br>n=3 | 11<br>n=1 | 8.4 (4.5; 15.7)<br>n=12 |
| Vax + BA.1 | 2.2 (1.7; 2.8)<br>n=32 | 1 (0.4; 2.9)<br>n=5 | 1 (0.7; 1.5)<br>n=10 |  | 1.3 (0.8; 2.1)<br>n=18 |
| Vax + BA.2 | 2.5 (1; 6)<br>n=6 | 3.8 (0.8; 17.4)<br>n=2 | 1.4 (0.5; 3.9)<br>n=2 |  | 1.8 (0.7; 4.1)<br>n=3 |
| WT conv | 19.5 (14.6; 26.1)<br>n=33 | 13.1 (7.3; 23.6)<br>n=7 | 4.1 (2; 8.1)<br>n=9 | 6.8 (2.5; 18.8)<br>n=4 | 10.6 (5.7; 19.7)<br>n=18 |
| Alpha conv | 27.4 (13; 57.7)<br>n=8 | 43.2 (21.1; 88.3)<br>n=7 | 10.2 (7.6; 13.8)<br>n=6 | 9.7 (4.2; 22.4)<br>n=4 | 14.9 (7.3; 30.6)<br>n=8 |
| Beta conv | 6.8 (3.5; 13.4)<br>n=6 | 11 (3.9; 31.5)<br>n=5 | 24.7 (15.2; 40.3)<br>n=5 | 18.1 (2.5; 131.8)<br>n=4 | 4.4 (1.4; 13.8)<br>n=6 |
| Gamma conv | 3.7 (1.2; 11.3)<br>n=4 | 4.2 (1.9; 9.2)<br>n=4 | 6.1 (3.6; 10.3)<br>n=4 | 15.1 (2; 112.7)<br>n=3 | 1.4 (0.9; 2.3)<br>n=4 |

Table 3: **BA.1 titer fold changes from variants per serum group (conv = convalescent).**
| Serum Group | WT | Alpha | Beta | Gamma | Delta |
| --- | --- | --- | --- | --- | --- |
| Delta conv | 6.3 (2.8; 14.1) | 6 (2; 18.1) | 2.5 (0.5; 11.8) | 3.2 (0.4; 28.9) | 36.7 (19.4; 69.4) |
|  | n=10 | n=6 | n=6 | n=4 | n=10 |
| BA.1 conv | 0.2 (0.1; 0.5) | 0.1 (0; 0.9) | 0.2 (0; 0.6) | 0.1 (0; 0.7) | 0.2 (0.1; 0.5) |
|  | n=12 | n=4 | n=7 | n=2 | n=10 |
| BA.2 conv | 1.4 (0.2; 8.5) | 0.8 | 0.8 | 0.4 | 1.3 (0.1; 30) |
|  | n=3 | n=1 | n=1 | n=1 | n=2 |

**Table 4:** Variant titer fold changes from WT (614D/G) per serum group (conv = convalescent). Mean fold drops were calculated from fold drops per study, not from GMT fold drops. Studies reporting fold drops but not GMTs result in a discrepancy between GMT based mean fold drop and individual study based mean fold drop (Table 2). The 95%CI is given in parentheses, the number of data points n in the next line. WT summarizes wild type-like strains (e.g: 614D, 614G)

| Serum Group | Alpha | Beta | Gamma | Delta |
| --- | --- | --- | --- | --- |
| 2x Vax | 1.3 (1; 1.7) | 4.6 (3.8; 5.5) | 2.6 (1.6; 4.2) | 2.7 (2.3; 3.3) |
|  | n=11 | n=44 | n=7 | n=50 |
| 3x Vax | 1 (0.7; 1.4) | 2.6 (2.1; 3.3) |  | 2.3 (1.9; 2.6) |
|  | n=9 | n=30 |  | n=33 |
| Inf + Vax | 1.7 | 3.1 (2.2; 4.2) | 3.3 | 1.8 (1.2; 2.8) |
|  | n=1 | n=12 | n=1 | n=16 |
| Vax + Inf | 1.2 (0; 504.7) | 5 (1.3; 18.9) | 3.2 | 1.9 (1.2; 2.9) |
|  | n=2 | n=3 | n=1 | n=10 |
| Vax + BA.1 | 1.1 (0.5; 2.3) | 2.2 (1.4; 3.7) |  | 1.8 (1.3; 2.5) |
|  | n=5 | n=10 |  | n=15 |

Table 4: **Variant titer fold changes from WT (614D/G) per serum group (conv = convalescent).**
| Serum Group | Alpha | Beta | Gamma | Delta |
| --- | --- | --- | --- | --- |
| Vax + BA.2 | 0.7 (0.7; 0.7) | 1.7 (1.7; 1.7) |  | 2.1 (0.1; 30) |
|  | n=2 | n=2 |  | n=3 |
| WT conv | 2.1 (1.5; 3) | 6.1 (3.7; 10) | 3.5 (1.2; 10.1) | 2.2 (1.7; 2.8) |
|  | n=7 | n=9 | n=4 | n=18 |
| Alpha conv | 0.6 (0.3; 1) | 2.7 (1; 7.4) | 2.6 (0.4; 15.4) | 1.8 (1.2; 2.8) |
|  | n=7 | n=6 | n=4 | n=8 |
| Beta conv | 0.8 (0.3; 2.3) | 0.3 (0.2; 0.8) | 0.4 (0.1; 3.2) | 1.5 (0.6; 3.9) |
|  | n=5 | n=5 | n=4 | n=6 |
| Gamma conv | 0.9 (0.5; 1.6) | 0.6 (0.3; 1.2) | 0.3 (0.1; 0.9) | 2.6 (0.9; 7.6) |
|  | n=4 | n=4 | n=3 | n=4 |
| Delta conv | 1.1 (0.7; 1.7) | 2.7 (1.4; 5) | 1.6 (0.3; 7.8) | 0.2 (0.1; 0.3) |
|  | n=6 | n=6 | n=4 | n=10 |
| BA.1 conv | 1.7 (1.1; 2.5) | 1.5 (0.8; 2.7) | 1 (0; 70) | 1.1 (0.8; 1.7) |
|  | n=4 | n=7 | n=2 | n=9 |
| BA.2 conv | 1.1 | 1.1 | 2.4 | 0.7 (0; 45.1) |
|  | n=1 | n=1 | n=1 | n=2 |

LOD censoring did not occur in the 3x Vax group where all WT and almost all BA.1 titers were detectable (Supplementary Fig. A3). Hence, the estimated average fold drop of 6.3x for this group is more reliable compared to the 2x Vax group and demonstrates the benefit of a third WT vaccination, which was much debated at the time of BA.1’s emergence. We investigated if the substantially lower fold drop in 3x Vax compared to 2x Vax is because higher titers in 3x Vax were underreported, either by laboratories not titrating to the endpoint, or because of a high-titer non-linearity in the assay by looking at titers and reported fold drops and found that the fold drop from WT to BA.1 was independent of titer magnitude against WT (Supplementary Fig. A3). A third vaccine dose also reduced fold changes more than non-Omicron breakthrough infections, with fold drops from WT of 15x and 9.4x in the Vax + Inf and Inf + Vax groups. Additionally, a third dose lifted titers against BA.1 above a level identified to be protective against symptomatic infection during WT circulation (Supplementary Fig. A3).

The effect of low WT titers on fold drops from WT to BA.1 is also seen in convaescent serum cohorts, despite the limited amount of data for most of these cohorts. While high titers against WT after infection with a WT-like variant resulted in mean fold drops from WT of 19.5x and 27.4x in the WT and Alpha conv (convalescent), respectively, mean fold drops in Beta and Delta conv were much lower, at 6.8x and 6.3x due to lower WT titers (Supplementary Fig. A3, Table 2-3). On the other hand, BA.1 fold drops relative to the infecting Beta and Delta variants were much higher compared to BA.1 fold drops from WT for the same groups: 24.7x and 36.7x (vs 6.8x and 6.3x) (Table 3). One possible mechanism is again a LOD effect reducing fold drops from WT in Beta and Delta convalescent serum groups (since these groups will have higher homologous titers compared to WT titers). Another possible mechanism could be neutralizing antibodies in Beta and Delta sera focusing on regions that are distinct from both WT and BA.1 and therefore, with respect to these sera, WT and BA.1 appear antigenically less distinct leading to lower titer differences between these antigens.

As expected, a (breakthrough) infection with either BA.1 or BA.2 substantially increased titers against BA.1, at times even above WT titer levels, and resulted in average fold drops smaller than 3x.

In general, we saw high variability of fold drop data within all serum groups in the whole dataset, likely owing to several factors such as age of participants, serum collection times, and different assays and cell types used to assess serum neutralization ability (Table 1). For the 3x Vax and WT conv groups, we found a statistically significant difference between fold changes measured using authentic virus (LV) and pseudovirus (PV), but while LV fold changes were significantly higher than PV in 3x Vax it was the other way around in WT conv (Supplementary Table A2). For reported GMTs however, we consistently found higher PV than LV GMTs, often at a significant level (Supplementary Tables A4-A5). Different serum collection times can affect fold drops either through a LOD mechanism as explained above or through higher cross-reactivity as demonstrated in Wilks. *et al.*[16]. Still, combining data from various sources gives quick and reliable results as shown in Fig. 2, and a public database where laboratories enter their results would greatly contribute to quick decision making, and further, assay refinement across laboratories.

### 2.4 Antigenic cartography of BA.1’s immune escape

As an additional way to evaluate the reliability of variable source data for immunological surveillance, we constructed antigenic maps from the collected data and compared them to single source antigenic maps. In an antigenic map, virus variants are positioned based on their antigenic properties inferred by fold drops in serum neutralization titers [11]. Variants that elicit similar titers in the same sera are positioned at small distances from each other, and vice versa. Antigenic maps are a key instrument for vaccine strain selection for influenza vaccines and have also been used to investigate SARS-CoV-2 vaccine strain updates [17].

Using data only from convalescent serum cohorts, we again found that data within one month of reporting produced a very stable result (Fig.3 a-b) (Supplementary Fig. A5-A10). Only Gamma’s position changed slightly when creating a map from all data (Fig. 3b). Already after one month of data reporting, the map captured BA.1’s complete escape from all pre-Omicron sera and variants. The early data map and the full data map were highly consistent with maps from single laboratories, both a map constructed using pseudovirus neutralization data [16] (Fig. 3c) and one using authentic virus [18] (Fig. 3d). Maps constructed using only authentic virus or pseudovirus neutralization data resulted in very similar variant positions for variants with sufficient titrations in the different serum groups (Supplementary Fig. A5).

**Fig. 3.**
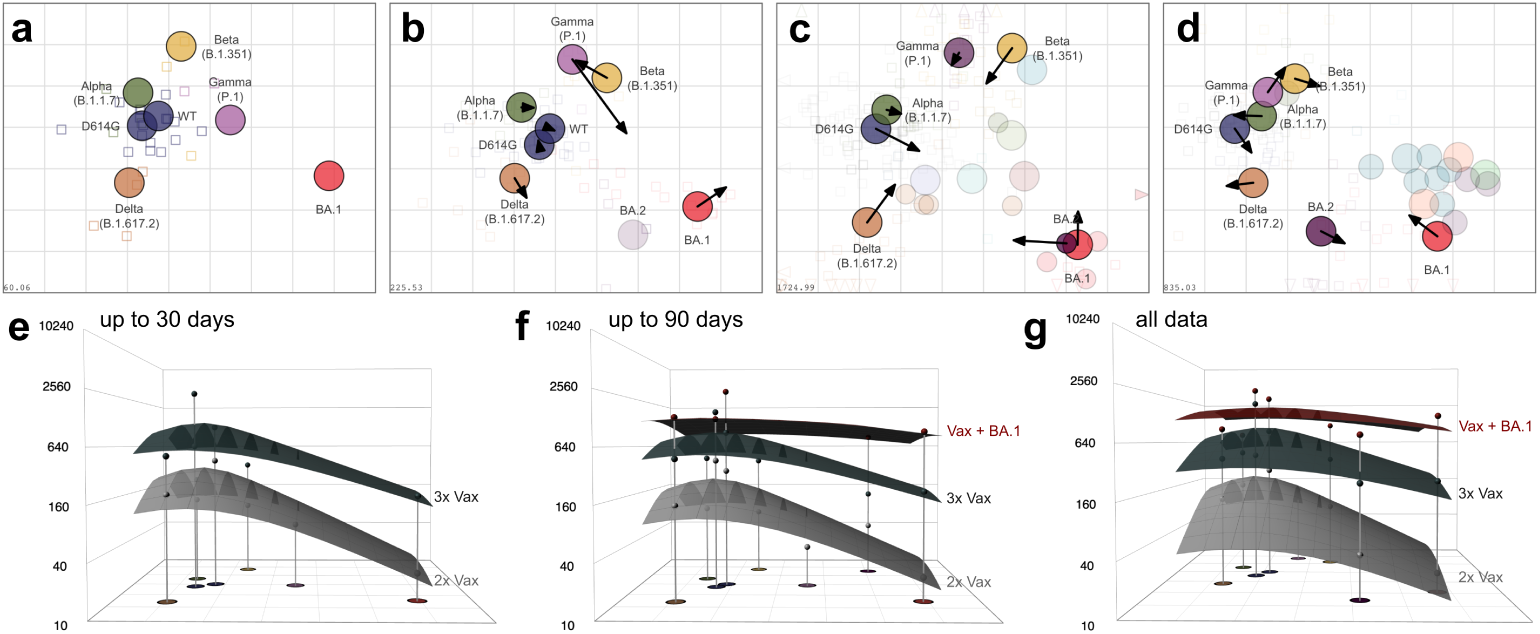
Antigenic cartography. **a)** An antigenic map from convalescent data up to 1 month post first available data was constructed. Variants are shown as labeled, colored circles and sera are shown as open squares with the color matching the infecting variant. The x- and y-axes correspond to relative antigenic distances, each grid line reflects an additional 2-fold dilution in the neutralization assay. b) An antigenic map constructed from all convalescent data with arrows pointing to the variants’ positions in a. c) Comparison of the map by Wilks et al. [16] to the full data map in b and d) the map by Roessler et al. [18] to the full data map in b. Arrows point to the variants’ positions in the respective map. The numbers on the bottom left corner show the stress of the map. e-f) GMT antibody landscapes for the 2x Vax (grey), 3x Vax (dark grey) and Vax + BA.1 (red) serum groups are shown, subset to early data up to 1 month post first report (e), medium data up to 90 days post first report (f) and all data (g). GMTs against variants are indicated by impulses

Whereas antigenic maps show individual sera as points based on their reactivity, antibody landscapes show the distribution of neutralization titers for individual sera against multiple strains as a surface in a third dimension above an antigenic map [19]. To illustrate the effect of a third vaccination or breakthrough infection we constructed antibody landscapes for the 2x Vax, 3x Vax and Vax + BA.1 serum groups, again for different end points (Fig. 3e-g). As before, we found that combining data from different sources gives stable representations of population immunity early after variant emergence. Moreover, we found a much heightened and broadened neutralizing response after, firstly, a third Wu-1 dose compared to only two doses, and secondly, BA.1 break-through compared to a third Wu-1 exposure. A third dose was necessary to lift BA.1 titers to a somewhat protective level compared to two doses (Supplementary Fig. A3).

### 2.5 Increased cross-neutralization over time since exposure

We finally sought to investigate if the collected data could be used to investigate biological phenomena previously described in single datasets. Wilks *et al.* [16] found that variant cross-neutralization increased over time after the second vaccination, and that cross-reactive antibodies were recalled after the third dose rather than induced by the third dose, indicating ongoing affinity maturation after the second dose. Following their approach, we constructed antibody landscapes for serum cohorts by time since exposure when the information was available. We binned individual sera by 2 weeks, 1, 3, 6, 9 and 12 months post exposure and fitted the landscape slope for these binned cohorts, with a smaller slope indicative of more cross-neutralization across antigenic space (Fig. 4).

**Fig. 4.**
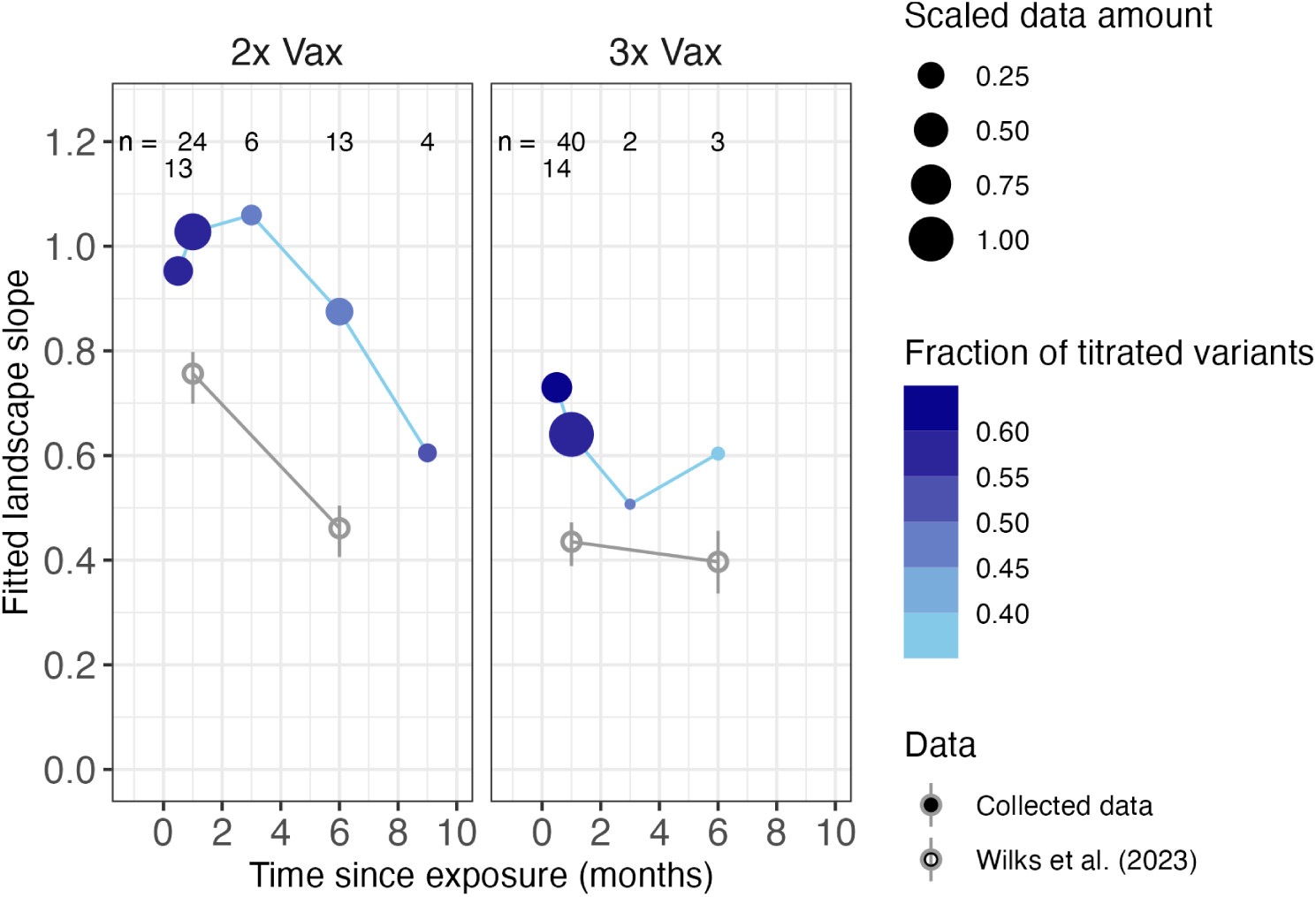
Antibody landscape slopes since exposure. Antibody landscapes were fitted to sera binned by their time since exposure for the 2x and 3x Vax cohorts. The time since exposure is given on the x-axis, the y-axis shows the fitted landscape slope. A low slope is indicative of broad cross-neutralization. The numbers in the top row indicate the number of sera that contributed to each landscape. The color of the point shows the fraction of variant titer measurements that contributed to the landscape fit, with a darker color indicating a more complete data set and better geometrical information for individual sera. The dot size gives the amount of data (n*fraction of titrated variants) per point scaled by the maximum data amount across serum cohorts. A detailed description is given n the methods section. The slopes reported by Wilks et al. [16] are shown in grey and not scaled by data or fraction of titrated variants

We found an overall trend of lower slopes as time since exposure increased, indicating higher cross-neutralization. In general, the data with information since exposure was scarce and binning resulted in additional inaccuracies, explaining incoherences of individual data points. Still, the slope value 9 months after the 2^nd^ dose is remarkably similar to the slope values half a month and one month after the 3^rd^ dose, supporting the hypothesis by Wilks *et al.* [16] that a third dose recalls broadened immunity rather than induces substantial broadening.

## 3 Discussion

Quick actions in response to the emergence of pandemic-causing pathogens are essential to minimize the disease’s toll on human lives and the global economy. When Omicron BA.1 started to circulate in late 2021, its accumulation of known escape substitutions in the spike protein [20] raised the concern that the Wu-1 targeted vaccine could be inefficient at protecting against BA.1, and that another lockdown might be necessary to prevent numerous hospitalizations and deaths. To provide decision makers and scientific advisory boards, such as the US NIH NAID SAVE (SARS-CoV-2 Assessment of Viral Evolution), the UKHSA SAGE (UK Health Security Agency Scientific Advisory Group for Emergencies) and the WHO, with the necessary information to recommend measures, we summarized publicly available data from preprints and publications from December 9, 2021 to August 14, 2022. During this time period, continuously updated versions of the here presented data were repeatedly presented and contributed to official advisory documents [2, 8–10]. Here, we presented a more extensive analysis of this data and demonstrated the immense utility of rapidly, and readily, available antigenic data for public health and pandemic responses.

In response to one of the most urgent questions in late 2021, whether the recommendation of a third Wu-1 vaccine applies during BA.1 circulation, some definitive statements could be made from the aggregated neutralization data. Sera from individuals who had been vaccinated twice or infected with WT once showed generally more than an 18x fold drop of titers from WT, whereas people who had been vaccinated three times showed average fold drops of 6.3x (Table 3). Moreover, a third dose raised titers above a level reported to be protective against WT symptomatic infection (Supplementary Fig. A3). The reduced titer drop in triple vaccinated individuals appears to be real and not an artifact of the assay’s limit of detection, as fold drops were consistent over a range of titers. Censored titers below an assay’s detection threshold can result in a deflation of fold drops when titers against the reference antigen are low. The mean fold drop for the 2x Vax group, for example, is likely substantially greater than our numeric estimate.

Remarkably, though, is that the above information was already in the public domain two weeks after BA.1’s emergence and remained largely stable thereafter (Fig. 2). Already after two weeks or ten random studies (Supplementary Fig. A2), we could dentify the benefit of a third dose, albeit with less certainty. We show that even summary data, such as the geometric mean titers, contain immensely valuable information when combined from different sources. This real-world information could help massively in designing vaccination strategies and clinical trials, saving time and financial resources by more targeted design. As an example, antigenic cartography was used in the design of the COVAIL trial to investigate SARS-CoV-2 vaccine strain updates [17]. Antigenic cartography from the collected data did not only give a robust representation of antigenic relationships between pre-Omicron and BA.1 and BA.2 variants, but also showed the advantage of distant exposures through BA.1 breakthrough infection in broadening immunity (Fig. 3), which was described in independent studies and the COVAIL trial [16, 17, 21].

Further, this analysis demonstrated that some of the phenomena observed in individual, more controlled, studies can also be seen in less controlled collated data. Wilks *et al.* [16] described an increase in cross-reactivity after the second vaccination over time, and the collated data corroborate this (Fig. 4). The lower fold drops and higher titers after a third vaccine dose could be the result of ongoing affinity maturation in germinal centers after a second vaccination, and a recall of affinity matured B cells upon the third dose. Kim *et al.* [22] reported the persistence of germinal centers for more than six months after mRNA vaccination, and found that antibodies derived from plasma cells with high levels of somatic hypermutation exhibited higher neutralization capacity against the D614G variant. Sokal *et al.* [23] found that memory B cell-derived monoclonal antibodies of vaccinated individuals could maintain binding capacity to the Beta variant during affinity maturation. This, in combination with broader antibody landscapes after the third dose, suggests that repeated vaccination with the original spike protein has the potential to boost cross-reactive immunity across antigenic space.

Finally, data from different sources provide valuable information for assay standardization. We found a statistically significant difference between pseudovirus- and authentic virus-assessed fold drops from WT to BA.1 in the WT conv and 3x Vax group. The collected data was generated using a variety of pseudotypes and cells, with different ACE2 and TMPRSS2 expression patterns. SARS-CoV-2 evolved in ways that elicited differential usage of TMPRSS2 for cell entry, resulting in different neutralization titers depending on TMPRSS2 expression of target cells [24–26]. We were limited by the amount of data and could not stratify by surface receptor expression to test this further. In general, the greatest limitation of aggregating independent datasets is the imited amount of data, the potential of misclassification due to human error, or biased data. In fact, the majority of subjects in the studies were female (Supplementary Fig. A11) and over a third of all studies came from the US; together with studies from Germany, they made up half of our dataset (Supplementary Fig. A12). Studies from countries from the Global South, and hence people, were massively underrepresented, highlighting research funding and access inequalities. Still, we argue that the trade-off between rapidly available but less-controlled data can be overcome when enough data s used, and has a clear benefit for quick and early pandemic responses. Additionally, an easily accessible database with uniform format and metadata on assay, serum, and localization could greatly contribute to answering questions about SARS-CoV-2 variant specificities for assay standardization, global immunity and public health focus. Tao et al. [27] showed the benefit of combining public data to assess BA.1’s and BA.2’s susceptibility to therapeutic monoclonal antibodies. In line with the proposed World Serology Bank by Metcalf *et al.* [28], we advocate for a public antigenic database, similar to sequence databases like GenBank and GisAid [29], where researchers upload virus neutralization and binding data at the time of uploading preprints to an online arXiv. ImmPort [30] contains antibody neutralization data inked with assay protocols and publications, but, firstly, the variety of data formats and, secondly, the time period from data generation to publication and upload, make t less suitable for quick pandemic responses. Alternatively, AI based tools have been employed to create a ‘living evidence map’ of COVID-19 [31], or extract existential risk information from publications [32]. A similar approach could be used to develop and maintain a ‘living antigenic database for SARS-CoV-2’. The development and employment of such technology could easily be applied to other pathogens. Climate change and invasion of humans into animal habitats will only lead to an increase of animal-to-human pathogen transmission; Together with the increase in global travel, global pandemics will only become more rather than less likely. We here make the case that an easily accessible public antigenic database, containing neutralization, binding, assay and serum data, would be immensely useful for pandemic preparedness and response.

## 4 Methods

### 4.1 Data collection

Omicron neutralization data from publicly available preprints (bioRxiv, medRxiv), reports or tweets were collected and metadata regarding serum type (infection history, time since exposure) and assay type (authentic virus, pseudovirus type, cell type) was extracted when available/found in the manuscript. The collected data was reported between 2021/12/08 and 2022/08/14. Mean neutralization fold drops and/or geometric mean neutralization titers were extracted as stated in the respective study’s manuscript. No individual sample data was used, when individual repeat data and the mean was reported we used the mean. Uncertainty of fold change reports was designated by “*>*” in case of *>*= 50% of samples below the assay’s limit of detection (LOD) of “*>>*” in case of *>*= 80%. For geometric mean titers (GMTs) the uncertainty was reported with “*<*” or “*<<*”, respectively. When no information on the number of detectable samples was given in the manuscript, the uncertainty was assigned by counting from the available data to the extent visual inspection allowed. A full list of all studies considered is shown in Table 1, detailed metadata in the publicly accessible excel sheet [3].

### 4.2 Serum group categorization

Serum panels were categorized by their infection or vaccination history into different serum groups. In the “2x Vax” group (n=48) we included double vaccinated individuals, independent of vaccine type, and single dose Johnson & Johnson (J&J) vaccinated ndividuals, as a single J&J dose is the recommended vaccination regime. The “3x Vax” (n=59) group consisted of triple vaccinated sera, or sera that received a combination of J&J and mRNA vaccines. Individuals with pre-Omicron infection and then vaccination were categorized as “Inf + Vax” (n=24), or vaccination and pre-Omicron breakthrough infection as “Vax + Inf” (n=13). Breakthrough infection with BA.1 was specified as “Vax + BA.1” (n=29) or “Vax + BA.2” (n=6). Convalescent “conv” sera were categorized by the infecting SARS-CoV-2 variant (n(WT)=29, n(Alpha)=8, n(Beta)=6, n(Gamma)=4, n(Delta)=10, n(BA.1)=15, n(BA.2)=4).

### 4.3 Geometric Mean Titer and fold drop calculation

Geometric Mean Titers (GMT) and mean titer fold drops to a reference antigen were used as stated in each study. In case of thresholded GMTs directly available or extracted from the manuscript, the GMT estimates were set to half the value of the imit of detection (Table 2). Fold drops were calculated by dividing the reference antigen GMT by the variant GMT. Omicron GMTs were obtained by applying the fold drop from wild type to the wild type GMT. Mean fold drops and 95% confidence intervals as reported in Tables 3-5 were obtained by calculating the mean of reported fold drops in each study using Rmisc’s CI() function (v 1.5.1) [33]. Some studies reported fold drops but not titers or variant titers resulting in discrepancies between individual study based mean fold drops and GMT based fold drops. The cumulative mean fold changes were calculated per day.

### 4.4 Antigenic cartography and antibody landscape slopes since exposure

Antigenic cartography was performed as previously described [16, 18, 34]. Antigenic maps were constructed in R [35] version 4.2.2 using the Racmacs [36] package (v 1.1.35) in 2 dimensions with 1000 optimizations using only convalescent sera. Map diagnostics were performed as previously described and can be found in the supplementary material (Supplementary Fig. A6-A10). Antibody landscapes were constructed as previously described using the ablandscapes [37] R package (v 1.1.0), ablandscape.fit() function with method=”cone”. This function fits a single cone surface to neutralization titers above an antigenic map. The cone coordinates and apex height is fitted per individual serum, the cone slope is optimized per serum group and quantifies cross-neutralization. A combined slope per serum group reduces overfitting for sera with few, sometimes only two, variant titrations. More variant titrations per serum improve the resolution of the plane across the antigenic map as more geometric information is given to fit the surface. The fraction of titrated variants quantifies this geometric information by dividing the number of measurements by the number of possible measurements (number of map antigens x number of sera). A steep slope indicates poor cross-neutralization. The data was subset to 30 or 90 days since first reported data to construct subset maps and landscapes. To calculate antibody landscape slopes since exposure, sera with information on time since exposure that were titrated against at least two variants were used. The sera were binned into 0.5, 1, 3, 6, 9, and 12 months since exposure. If the time since exposure was more than previous month + (next month – previous month)/2, the sera was assigned to the next month bin. To quantify the amount of data contributing to each slope, an additional measure of uncertainty, the number of sera n was multiplied with the fraction of available titer measurements, which is calculated by dividing the number of actual titrations by the number of possible measurements. The number of possible measurements is given by n(map variants) x n(serum group sera). To compare data availability across serum groups and times since exposure, the such calculated data amount was scaled by the maximum data amount across serum groups and times since exposure.

## Acknowledgements

Thanks and kudos to the laboratories listed here that rapidly generated data and put it into the public domain. We thank Poppy Roth for technical assistance.

## Data availability

The aggregate dataset is available as a publicly accessible google sheets document [3].

## Code availability

All code is publicly available in the manuscript’s GitHub repository (https://github.com/acorg/netzl et al2025.git). A citeable DOI will be generated upon manuscript acceptance.

## Author contributions

Conceptualization: All

Methodology: Antonia Netzl, Samuel H. Wilks, Sina Tureli, Derek J. Smith Software: Antonia Netzl, Sina Tureli, Eric LeGresley, Samuel H. Wilks

Validation: Antonia Netzl, Sina Tureli, Derek J. Smith

Formal analysis: Antonia Netzl, Sina Tureli

Investigation: Antonia Netzl, Sina Tureli, Eric LeGresley, Barbara Mühlemann, Derek J. Smith

Data curation: Antonia Netzl, Sina Tureli, Eric LeGresley

Writing - Original Draft: Antonia Netzl

Writing - Review and Editing: All

Visualization: Antonia Netzl

Supervision: Derek J. Smith

## Appendix A Supplementary Materials

### A.1 Supplementary Figures

Generated at the end of the document.

**Fig. A1.**
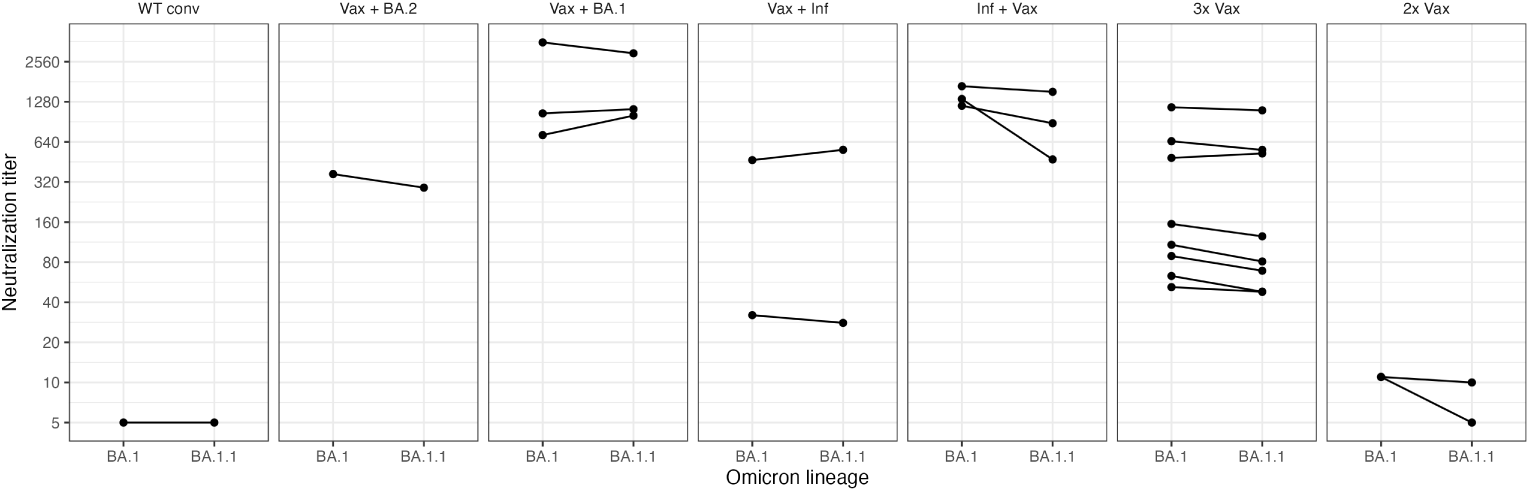
Neutralization titers of BA.1 and BA.1.1. Titers against the two lineages are shown or individual sera in which both lineages were titrated. Measurements from the same sera are connected by lines. The data is stratified by serum group

**Fig. A2.**
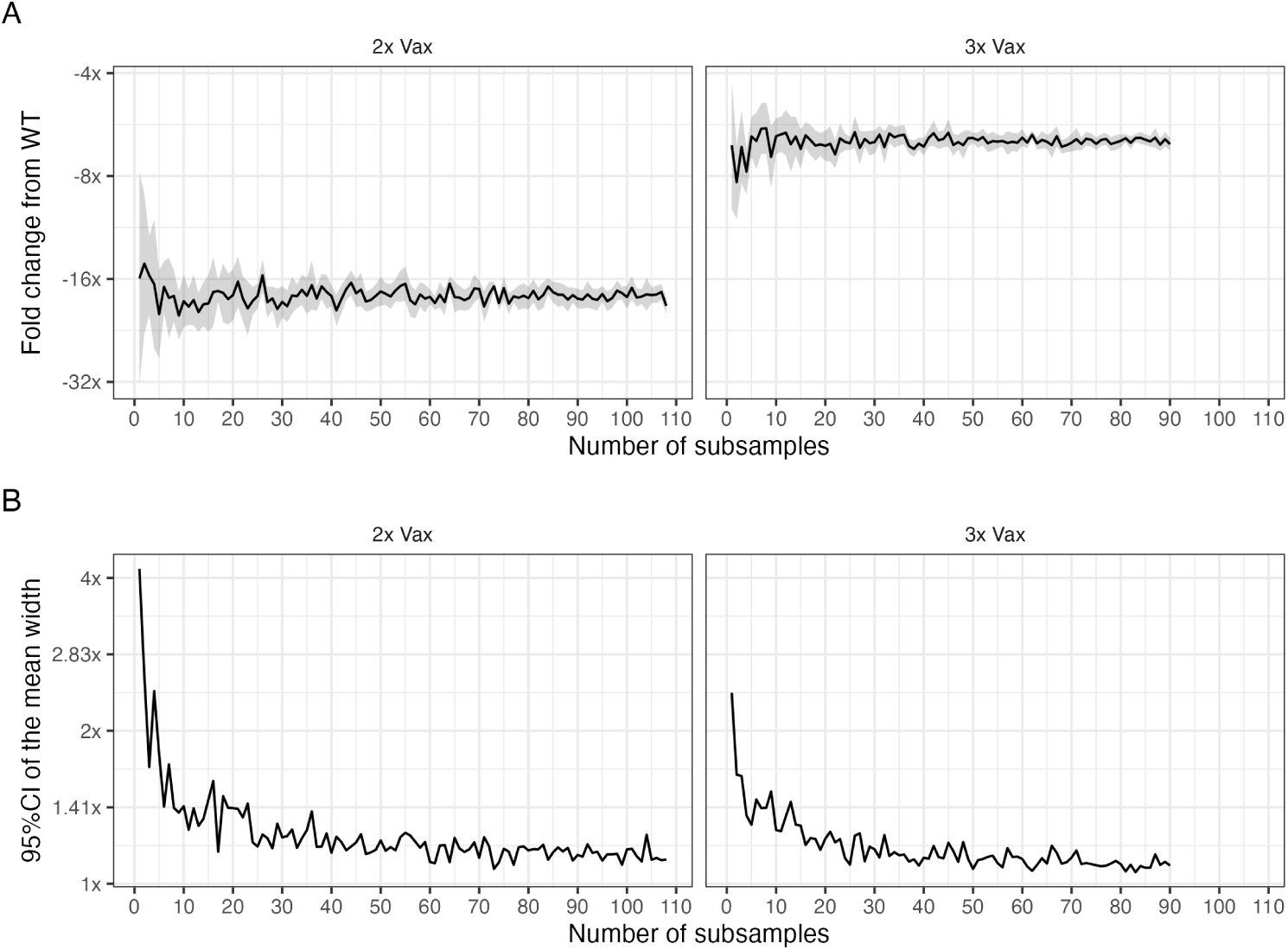
Bootstrapping the number of studies for BA.1 fold change assessment. **A)** The titer fold change from WT (614D/G) to BA.1 in the 2x and 3x Vax cohorts was calculated for n=1 to n=total number of studies per cohort by randomly subsampling n studies with replacement. For each n, the random selection was repeated 11 and 9 times in the 2x and 3x Vax cohort respectively, corresponding to 10% of the total number of studies per cohort. The geometric mean and its 95% confidence interval are shown as a solid black line with shaded area. **B)** The range of the such calculated 95%CI of the mean is shown across the number of subsampled studies for the 2x and 3x Vax cohorts (Upper - Lower 95%CI on the log2 scale, corresponding to Upper/Lower 95%CI on the linear scale)

**Fig. A3.**
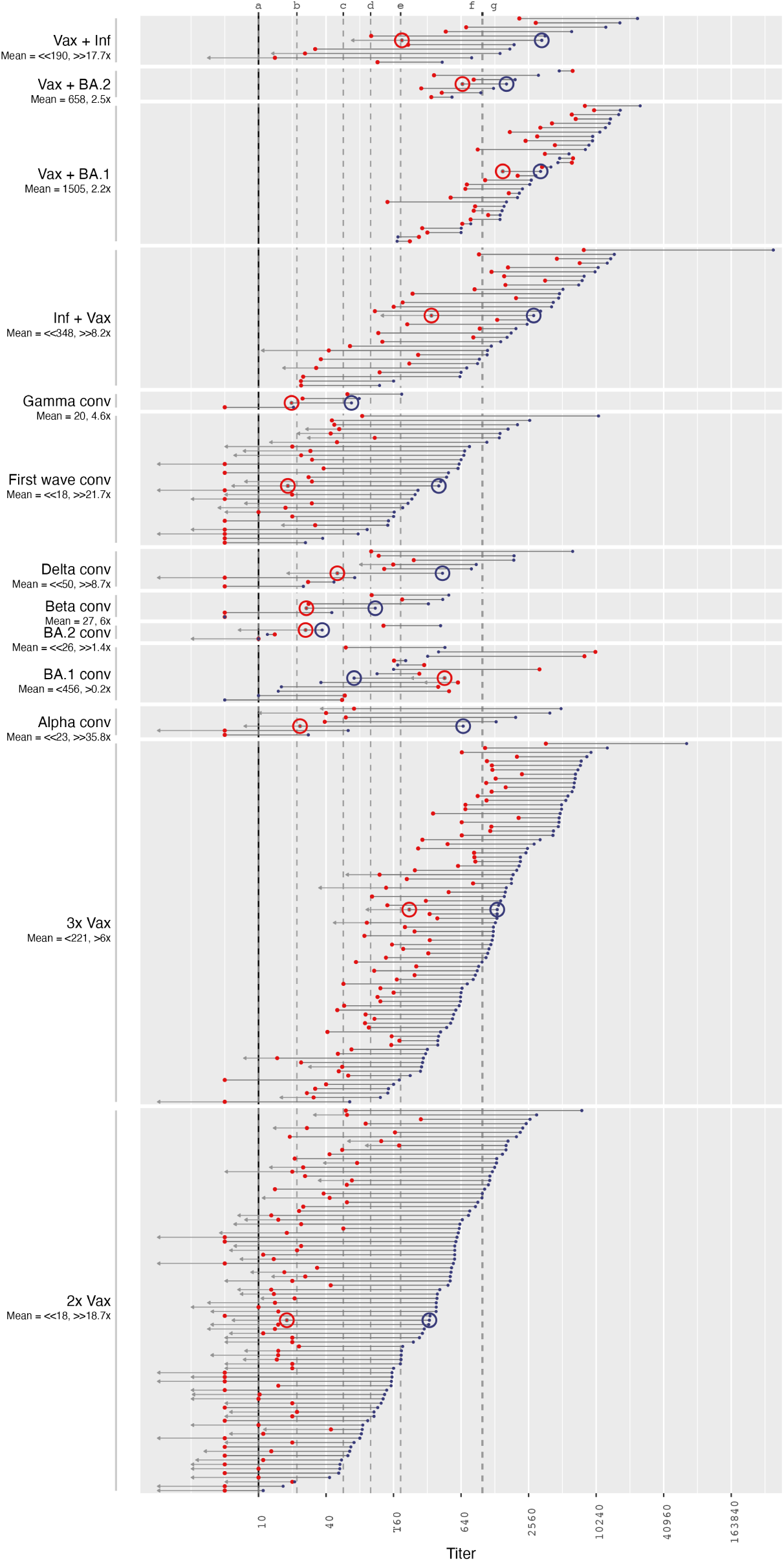
Omicron BA.1 and WT (614D/G) neutralization titers. GMT neutralization titers as reported by individual studies are shown, ordered by magnitude and stratified by serum group. Red dots indicate Omicron titers and small blue dots indicate WT titers, the big circles indicate GMTs per grouping. Below the serum group label, the BA.1 geometric mean titer (GMT) is given followed by the mean fold change from WT titers. The mean fold change from WT to BA.1 is indicated by the horizontal bar connecting WT and BA.1 titers. Arrows indicate uncertainties in the point estimate due to titers below the limit of detection (LOD)sdf of the assay. A short arrow (*>*/*<*) marks groups with more than half of BA.1 titers below the assay’s LOD, or conversely reference antigen titers at or lower than the LOD. Long arrows (*>>\<<*) mark groupswith more than approximately 80% of BA.1 titers below the LOD. Dashed lines mark thresholds of protection against symptomatic disease after vaccination with two doses of Moderna (a,d,f,g) [114] or AstraZeneca (b,c,e,f) [115] assessed by pseudovirus neutralization assay (a 78% VE, b 60% VE, c 70% VE, d 91% VE, e 80% VE, f 90% VE, g 96% VE)

**Fig. A4.**
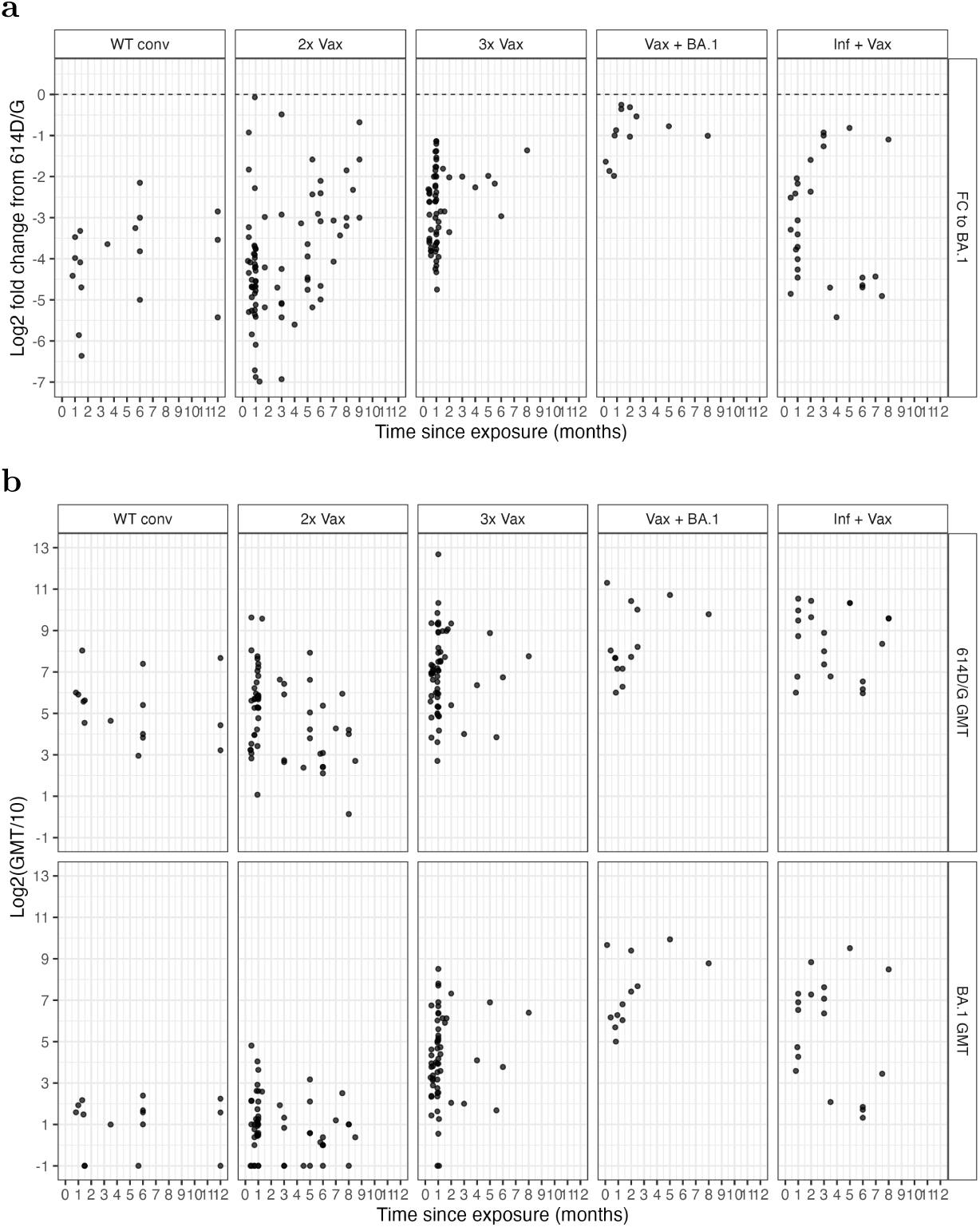
Individual studies by time since last exposure. **a)** 614D/G (WT) to BA.1 log2 fold changes by time since last exposure and **b)** geometric mean titers (GMTs) for 614D/G (WT) and BA.1 by time since last exposure in the individual studies which listed time since exposure information. The data is stratified by exposure history

**Fig. A5.**
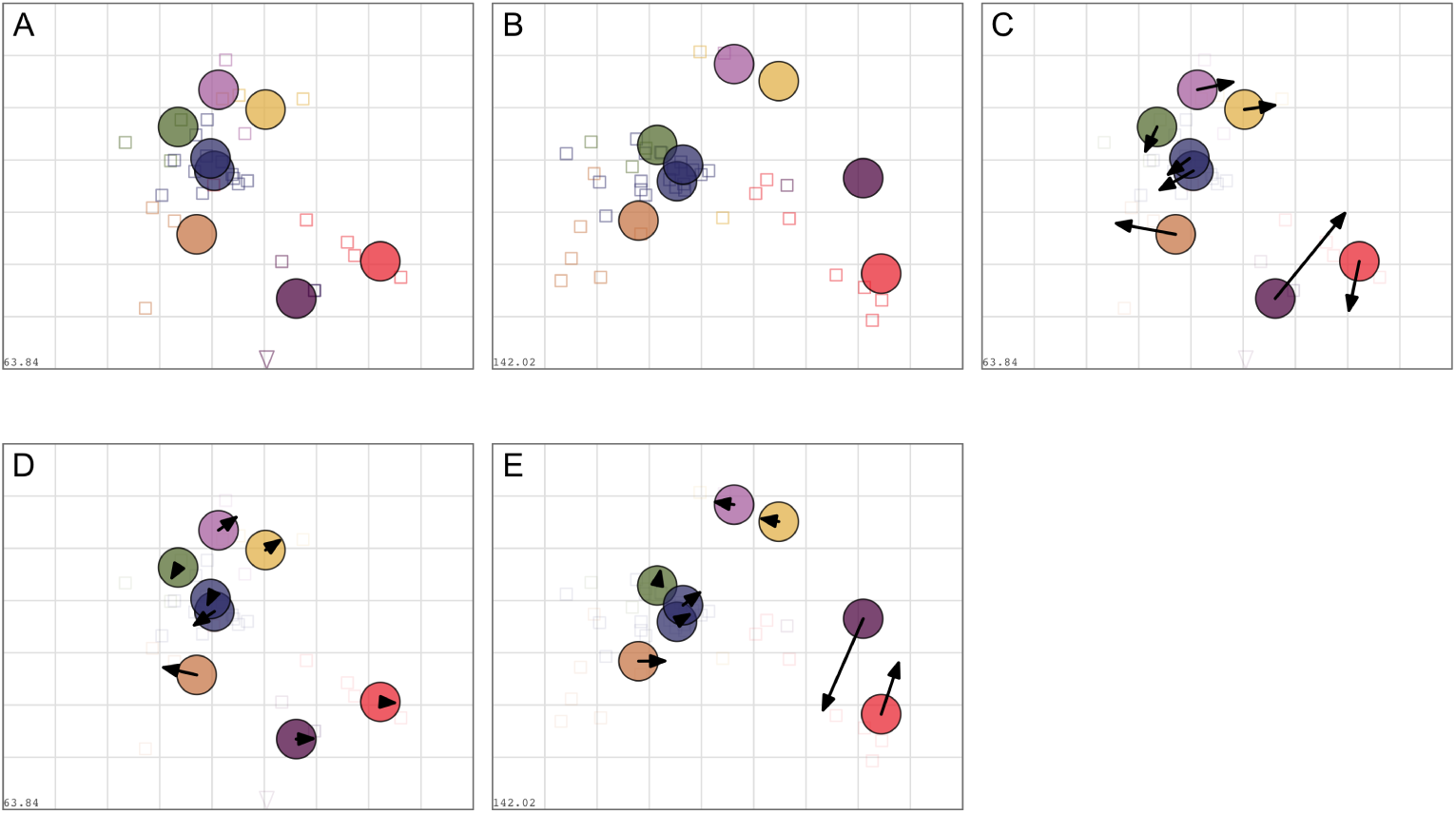
Antigenic maps from authentic virus and pseudovirus data. Antigenic maps were constructed as described in the methods section. **A)**A map was constructed using only data from authentic virus neutralization assays and **B)** using only data from pseudovirus neutralization assays. **C)** Shows a comparison of the two maps with arrows pointing from the variant position in map A to the variant position in map B. **D)**Shows the same comparison but with arrows pointing from A to the full data map shown in Fig. 3b. **E)** Shows a comparison of map B with arrows pointing to the ull data map shown in Fig. 3b

**Fig. A6.**
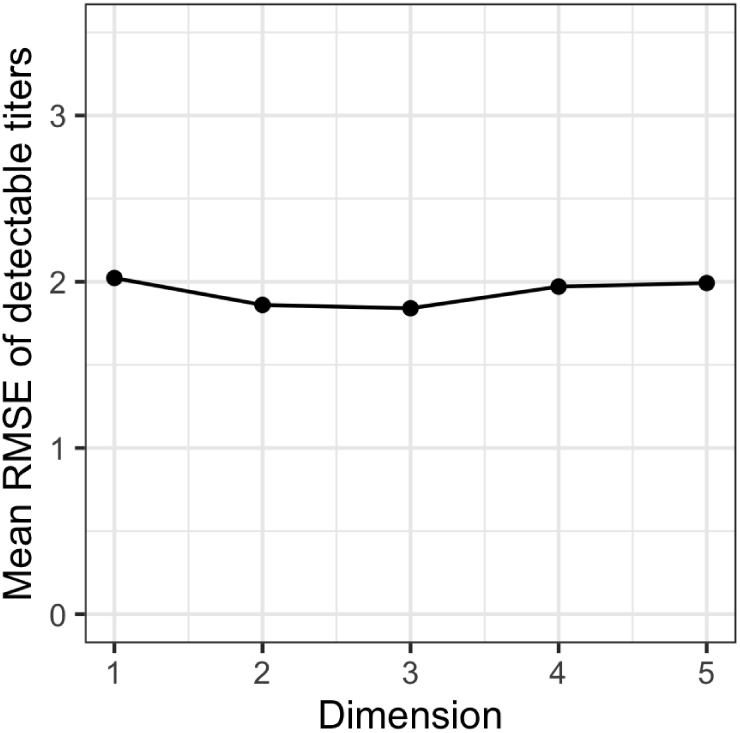
Dimensionality Test. Root Mean Squared Error (RMSE) between map and measured titers for detectable titers in 1 to 5 dimensions. Per dimension, 100 map replicates were constructed from 90% of measured titers with 1000 optimizations per replicate. The titers of the remaining 10% were predicted in each run and the RMSE calculated by comparing the predicted to the measured titers on the log_2_ scale

**Fig. A7.**
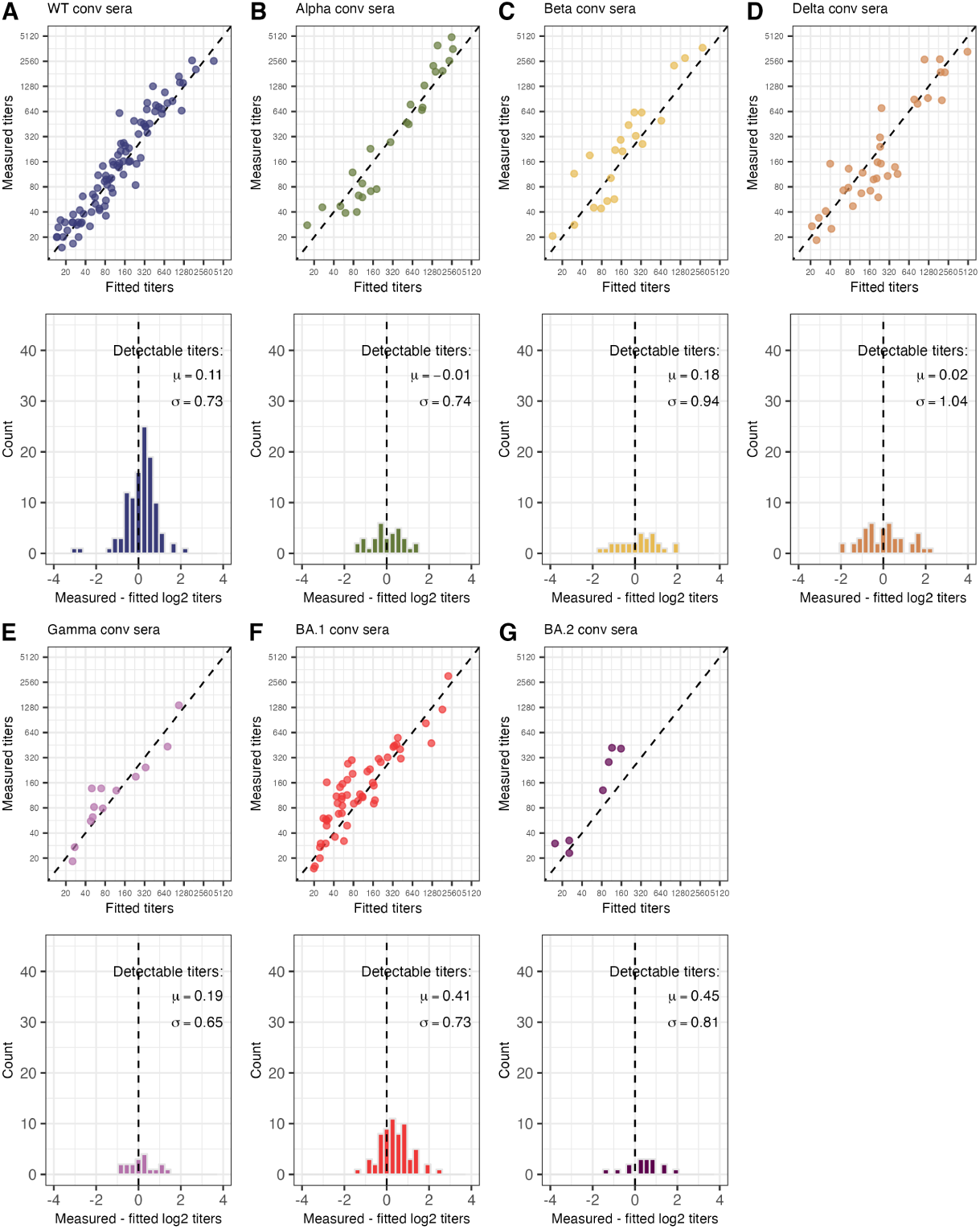
Goodness of map fit per serum group. The top panels show the correlation of detectable measured and fitted titers in the 2D map (Fig. 3b). Map distances were converted into og_2_ titers by subtracting the Euclidean distance for each serum-antigen pair from the maximum log_2_ titer of the specific serum. The bottom panels show the residuals of measured against fitted titers on the log_2_ scale, light grey marks pairs with the measured titer below the assay detection threshold. The mean and mean-centered standard deviation of differences between fitted and detectable measured titers are given in the legend of each bottom row panel. This was done for the serum groups used to construct the map: Infected with **A** WT (614D/G), **B** Alpha, **C** Beta, **D** Delta, **E** Gamma, **F** BA.1. and **G** BA.2

**Fig. A8.**
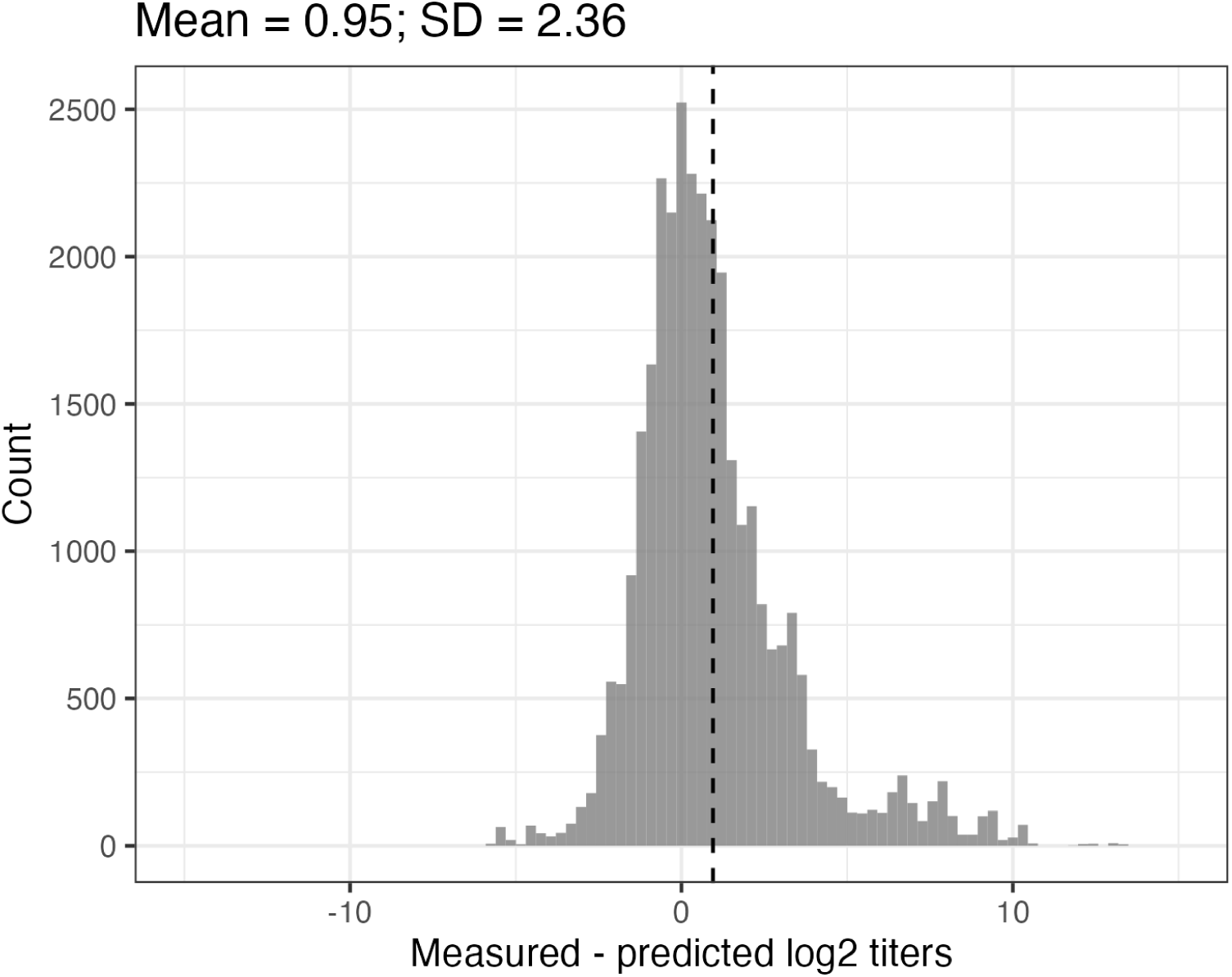
Map cross-validation residual titers. 1000 repeats with 1000 optimization runs each were performed with only 90% of measured titers used for map construction by artificially masking 10% of measurements. The missing log_2_ titers were predicted by subtracting the Euclidean map distance for each serum-antigen pair from the maximum log_2_ titer of the specific serum. The difference between predicted and detectable measured titers on the log_2_ scale was calculated, the mean is indicated by the dashed line. The mean and mean-centered standard deviation are given in the title

**Fig. A9.**
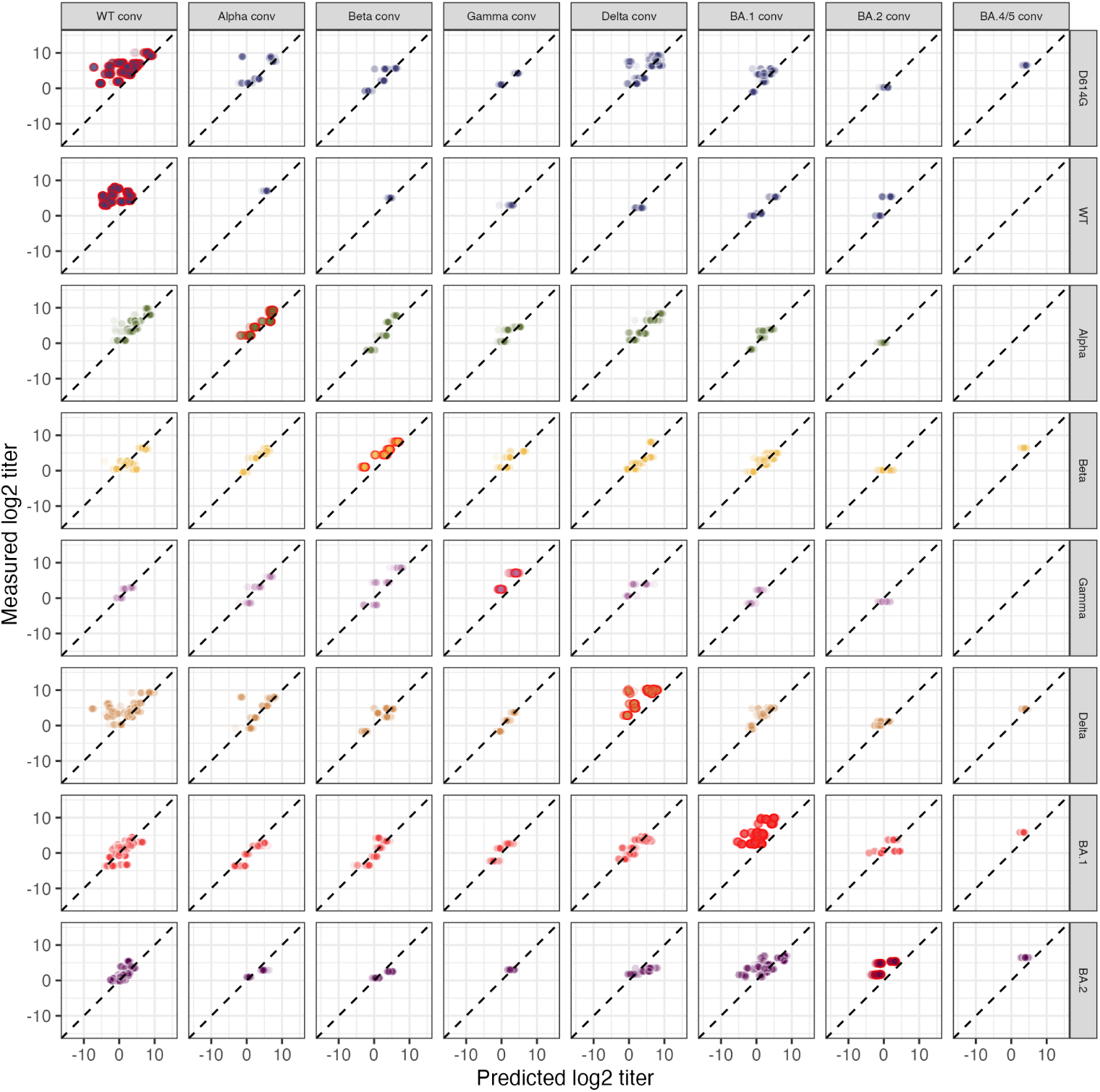
Predicted vs. measured titers. 1000 repeats with 1000 optimization runs each were performed with only 90% of measured titers used for map construction by artificially masking 10% of measurements. The missing log_2_ titers/10 were predicted by subtracting the Euclidean map distance or each serum-antigen pair from the maximum log_2_ titer/10 of the specific serum. Measured over predicted log_2_ titers/10 are shown per serum group and antigen variant. Virus variants are shown as rows, serum groups as columns. The points are coloured by variant and a red outline is added for the homologous serum-antigen pairs

**Fig. A10.**
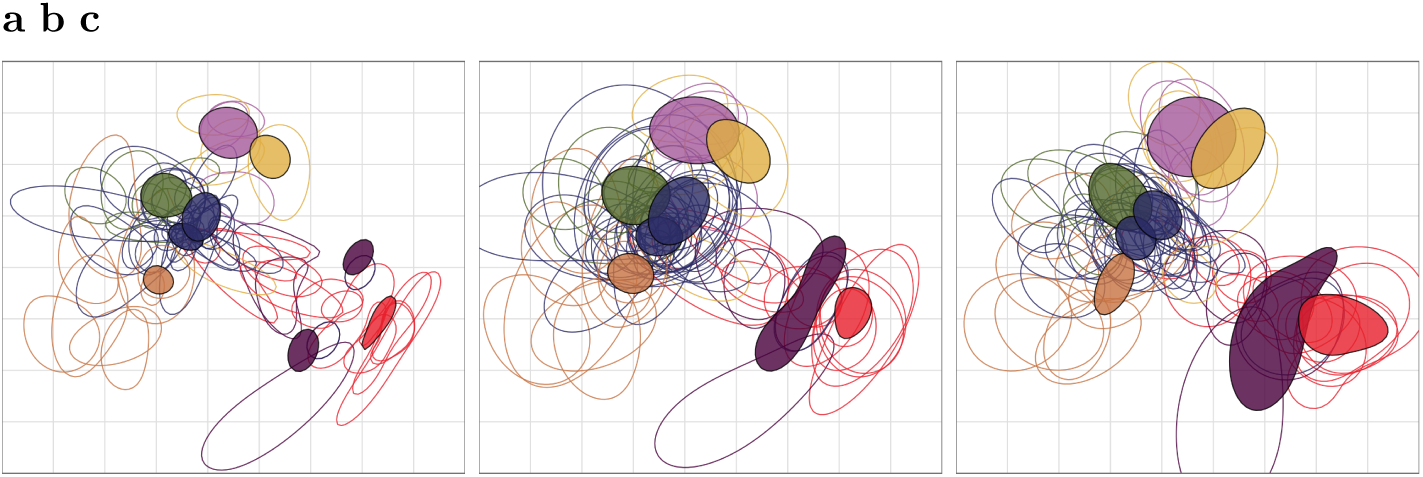
Positional resolution by bootstrapping. 500 bootstrap repeats were performed with 1000 optimizations per repeat and options as listed in the Methods section. In each repeat, **a)** different weights are assigned to sera and antigen reactivity of the titer table. The weights are drawn randomly from a Dirichlet distribution. **b)** Sera and antigens are randomly resampled with replacement, and **c)** normally distributed noise with sd = 0.7 is added to antigens and individual titer measurements. The colored regions mark 68% (one standard deviation) of the positional variation for each variant (filled shapes) and sera (open shapes). The colors correspond to the colors used in Fig. 3

**Fig. A11.**
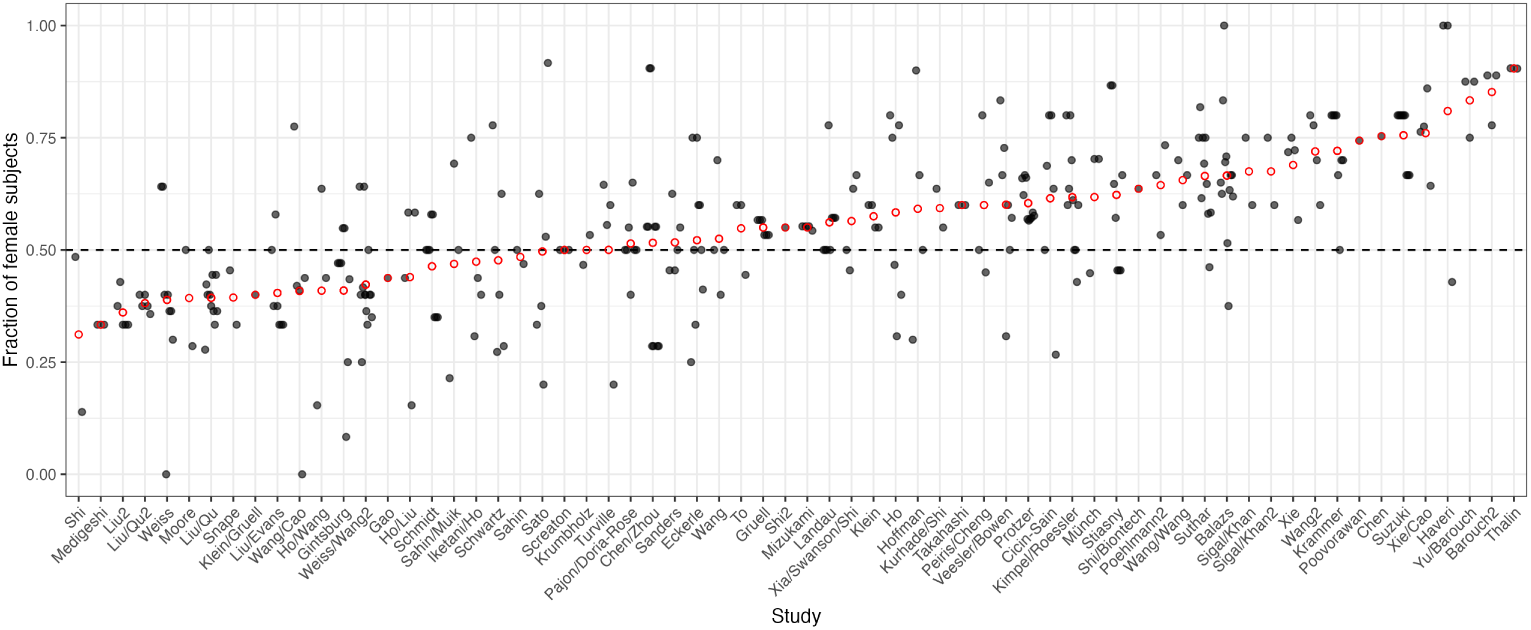
Fraction of female subjects per study. The fraction of females out of all study subjects was calculated for studies where sex metadata was found. Serum groups per study are shown as black dots with a jitter added on the x-axis to minimize overplotting, the mean fraction of females per study is given by the red, open circle. The dashed line indicates 50%. 29 studies reported data with some or all of the sex metadata missing, 66 studies reported some or all of the sex metadata

**Fig. A12.**
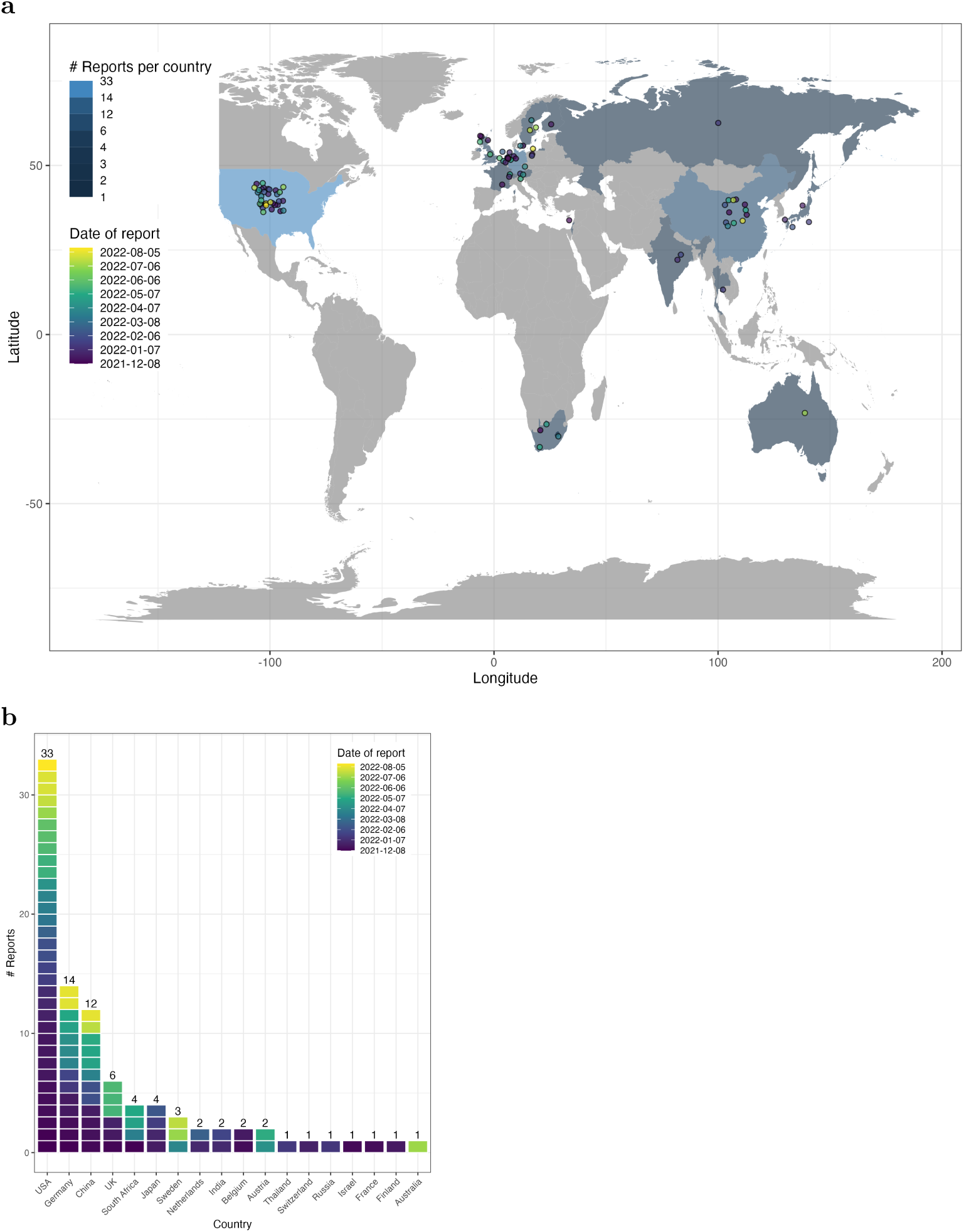
Studies by location and date. **a** Individual studies are located in the country of their corresponding authors’ institution as colored circles. The color corresponds to the date of online reporting, ranging from blue for early reports to yellow for late reports. The blue country color indicates the number of studies per country, increasing in brightness with increasing number of studies. Grey color indicates that no reports from these countries were used in the current study. **b**Shows the information in **a**in bar plot form, ordered by decreasing number of studies per country

### A.2 Supplementary Tables

**Table A1:**
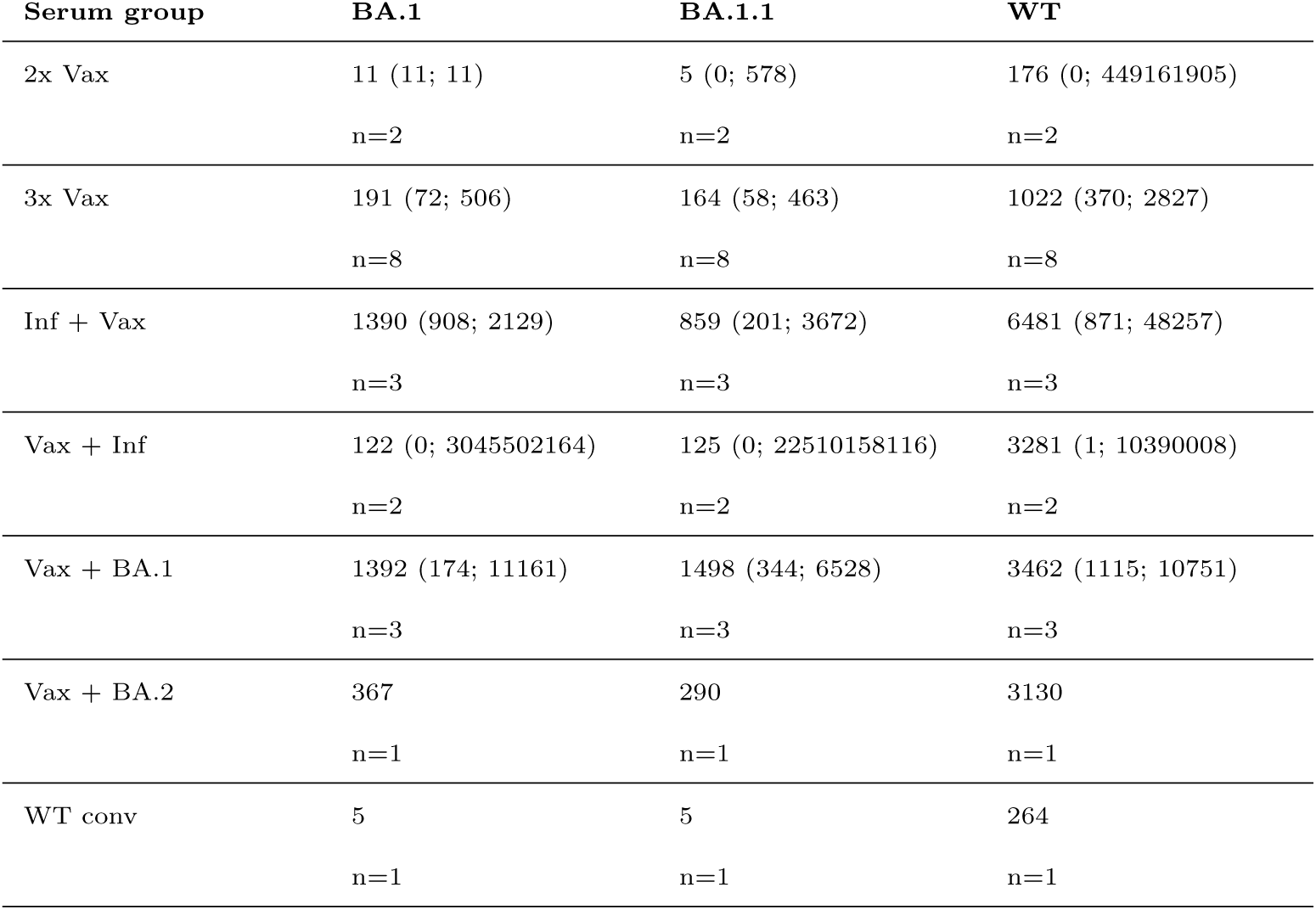
Geometric mean titer of BA.1 and BA.1.1. GMTs from individual sera that titrated both BA.1 and BA.1.1 are shown with 95% confidence intervals

**Table A2:** Assay-based statistical comparison of live virus vs pseudovirus fold changes from WT/D614G to BA.1. A t-test was performed to compare pseudovirus (PV) and live virus (LV) assessed fold changes in different serum groups. Normality was checked with a Shapiro-Wilk test and these serum groups were found to not differ significantly from a normal distribution. Results are ordered by increasing p-value. A statistic above 0 indicates higher values in PV than LV. The effect size was calculated with Cohen’s d for unequal variances and Hedge’s correction due to small sample sizes. All tests were performed with the rstatix package [116]

| Serum group | n(PV) | n(LV) | statistic | 95%CI (lower;upper) | df | p | Significance | Effect size | Magnitude |
| --- | --- | --- | --- | --- | --- | --- | --- | --- | --- |
| 3x Vax | 61 | 28 | -4.39 | -1.11;-0.42 | 51.4 | 0.00 | **** | -1.00 | large |
| WT conv | 19 | 14 | 2.99 | 0.35;1.83 | 30.9 | 0.01 | ** | 1.01 | large |
| 2x Vax | 69 | 39 | -1.58 | -0.98;0.11 | 88.1 | 0.12 | ns | -0.31 | small |
| Beta conv | 3 | 3 | -0.97 | -3.51;2.02 | 2.5 | 0.42 | ns | -0.64 | moderate |
| Vax + Inf | 10 | 5 | -0.84 | -2.31;1.1 | 6.9 | 0.43 | ns | -0.45 | small |
| Vax + BA.1 | 25 | 7 | -0.59 | -1.4;0.83 | 8.2 | 0.57 | ns | -0.26 | small |
| Delta conv | 7 | 3 | 0.47 | -2.14;3.12 | 5.2 | 0.66 | ns | 0.28 | small |

**Table A3:** Assay-based statistical comparison of live virus vs pseudovirus fold changes from WT/D614G to BA.1. A Wilcoxon-test was performed to compare pseudovirus (PV) and live virus (LV) assessed fold changes in different serum groups. Normality was checked with a Shapiro-Wilk test and these serum groups were found to differ significantly from a normal distribution. Results are ordered by increasing p-value. A statistic above 0 indicates higher values in PV than LV. The effect size was calculated with Wilcoxon effect size. All tests were performed with the rstatix package [116]

| Serum group | n(PV) | n(LV) | statistic | 95%CI (lower;upper) | p | Significance | Effect size | Magnitude |
| --- | --- | --- | --- | --- | --- | --- | --- | --- |
| Inf + Vax | 17 | 23 | 138.5 | -2.03;0.2 | 0.12 | ns | 0.25 | small |

Table A3: **Assay-based statistical comparison of live virus vs pseudovirus fold changes from WT/D614G to BA.1.**
| Serum group | n(PV) | n(LV) | statistic | 95%CI<br>(lower;upper) | p | Significance | Effect size | Magnitude |
| --- | --- | --- | --- | --- | --- | --- | --- | --- |
| BA.1 conv | 6 | 5 | 12.0 | - 6.99;2.9 | 0.65 | ns | 0.17 | small |
| Alpha conv | 5 | 3 | 9.0 | - 1.74;3.65 | 0.79 | ns | 0.16 | small |

**Table A4:**
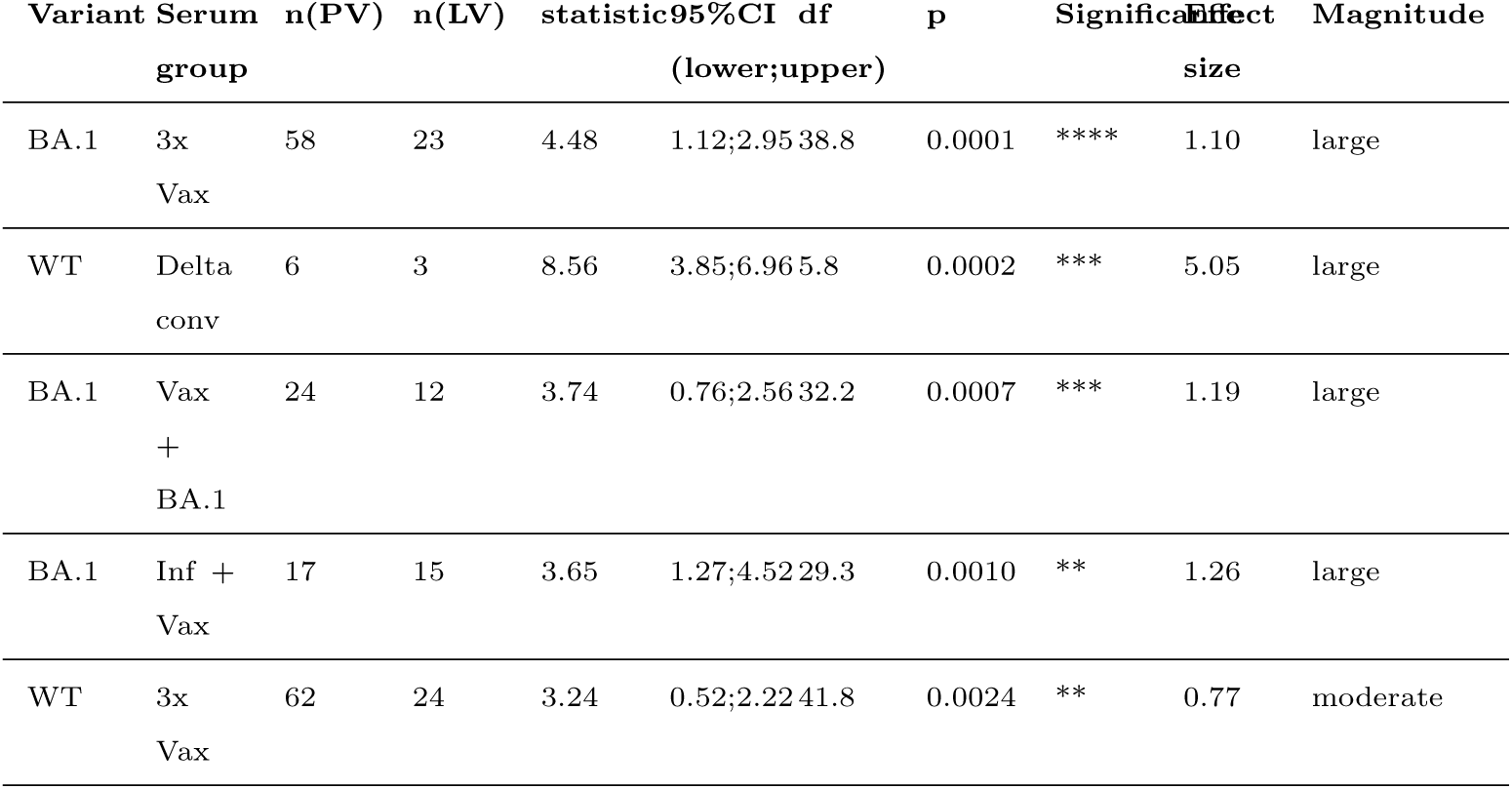
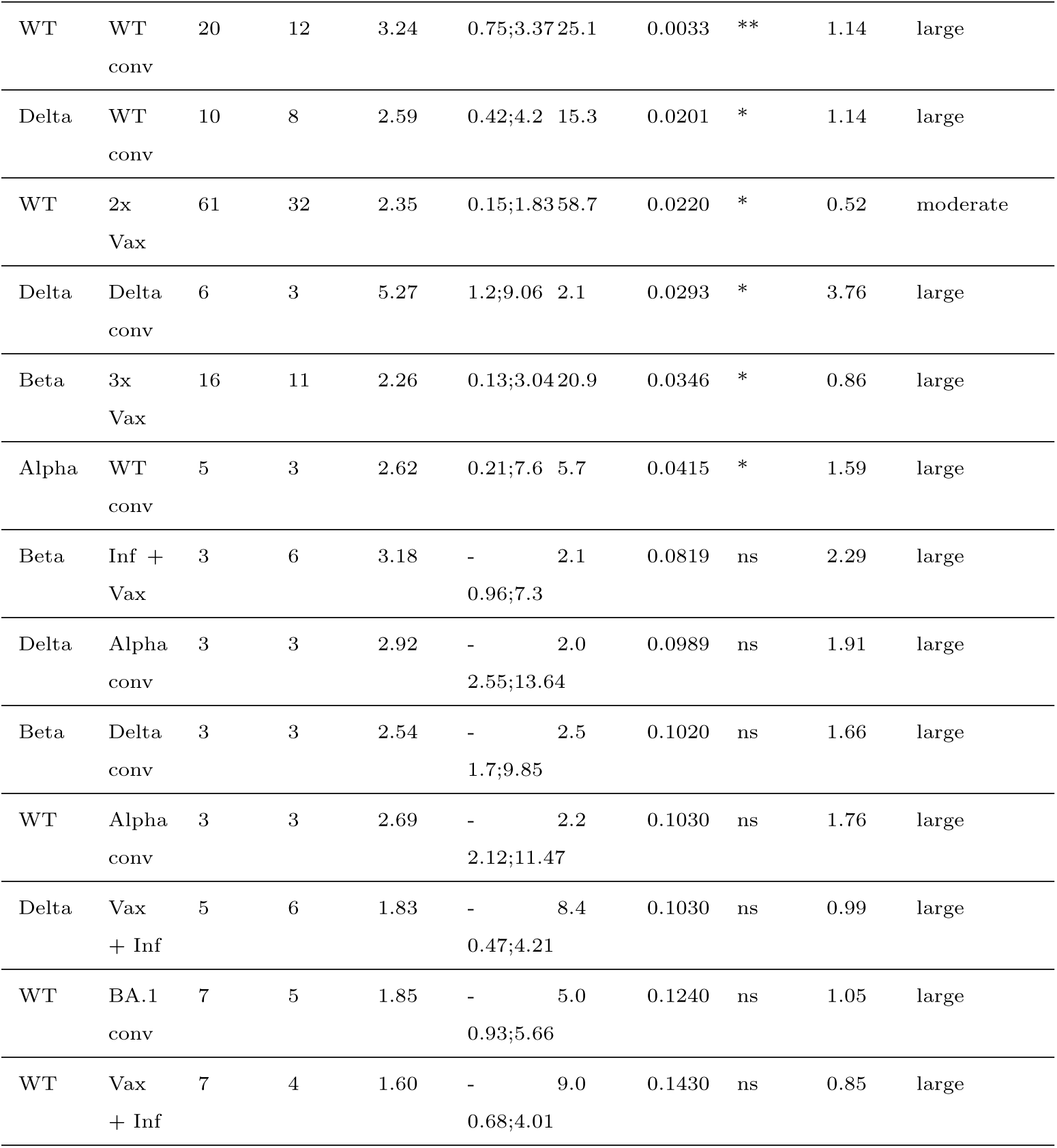
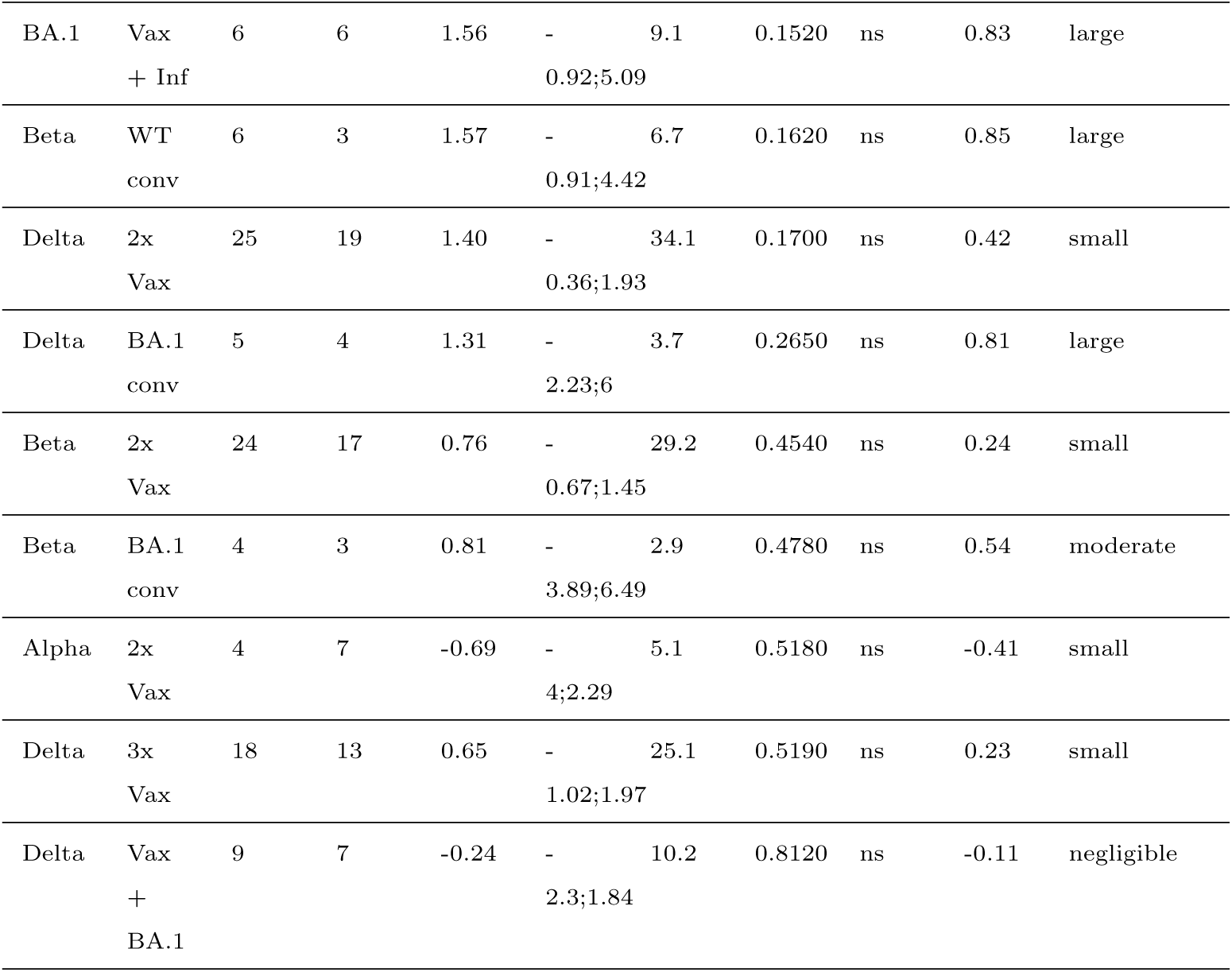
Assay-based statistical comparison of live virus vs pseudo virus variant GMTs. A t-test was performed to compare pseudovirus (PV) and live virus (LV) assessed Geometric Mean Titers (GMTs) in different serum groups. Normality was checked with a Shapiro-Wilk test and these serum groups were found to not differ significantly from a normal distribution. Results are ordered by increasing p-value. A statistic above 0 indicates higher values in PV than LV. The effect size was calculated with Cohen’s d for unequal variances and Hedge’s correction due to small sample sizes. All tests were performed with the rstatix package [116]

**Table A5:**
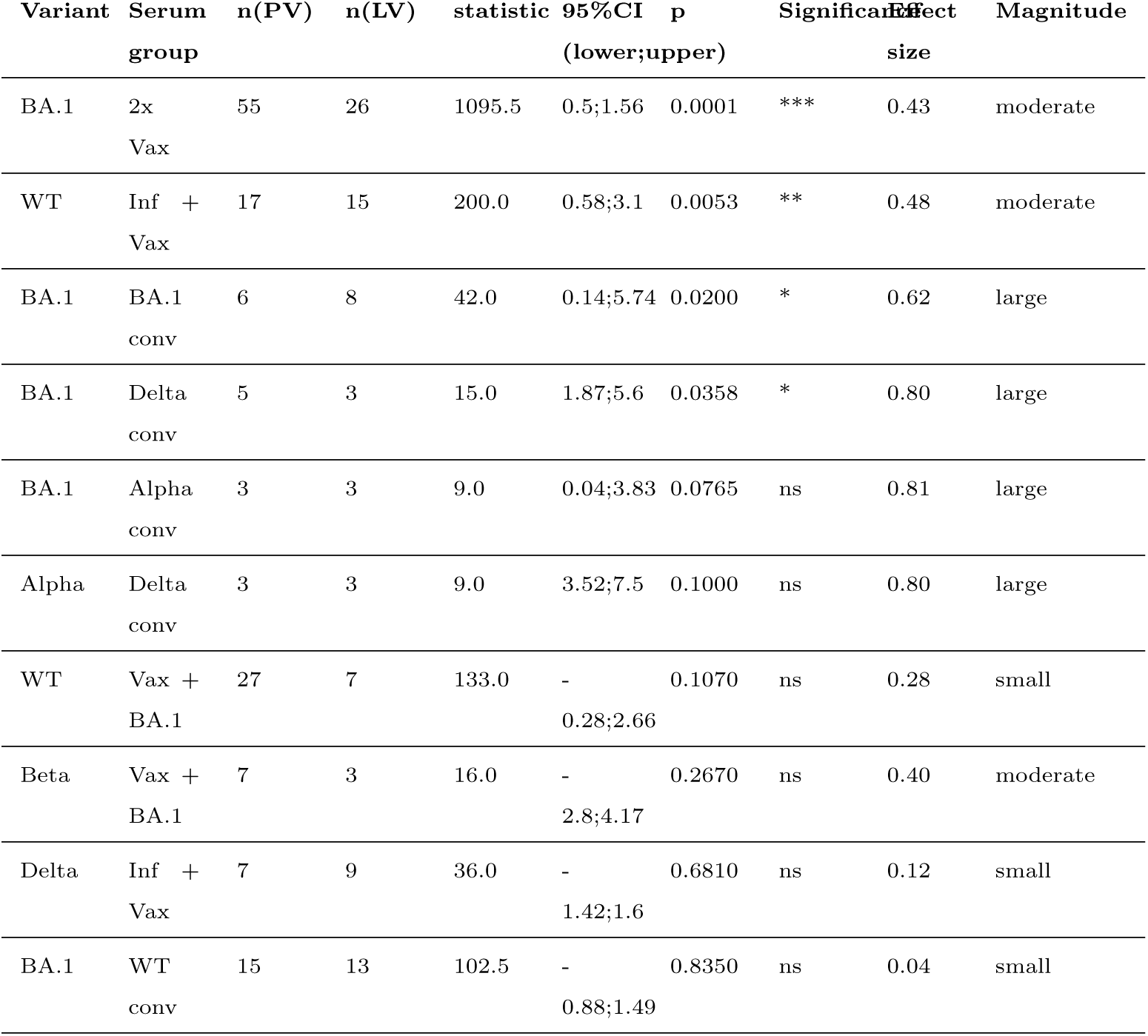
Assay-based statistical comparison of live virus vs pseudovirus variant GMTs. A Wilcoxon-test was performed to compare pseudovirus (PV) and live virus (LV) assessed Geometric Mean Titers (GMTs) in different serum groups. Normality was checked with a Shapiro-Wilk test and these serum groups were found to differ significantly from a normal distribution. Results are ordered by increasing p-value. A statistic above 0 indicates higher values in PV than LV. The effect size was calculated with Wilcoxon effect size. All tests were performed with the rstatix package [116]

